# Deaminase-assisted single-molecule and single-cell chromatin fiber sequencing

**DOI:** 10.1101/2024.11.06.622310

**Authors:** Elliott G. Swanson, Yizi Mao, Benjamin J. Mallory, Mitchell R. Vollger, Jane Ranchalis, Stephanie C. Bohaczuk, Nancy L. Parmalee, James T. Bennett, Andrew B. Stergachis

## Abstract

Gene regulation is mediated by the co-occupancy of numerous proteins along individual chromatin fibers. However, our tools for deeply profiling how proteins co-occupy individual fibers, especially at the single-cell level, remain limited. We present Deaminase-Assisted single-molecule chromatin Fiber sequencing (DAF-seq), which leverages a non-specific double-stranded DNA deaminase toxin A (SsDddA) to efficiently stencil protein occupancy along DNA molecules via selective deamination of accessible cytidines, which are preserved via C-to-T transitions upon DNA amplification. We demonstrate that DAF-seq enables ∼200,000-fold enrichment of target loci for single-molecule footprinting at near single-nucleotide resolution, enabling the precise delineation of the regulatory logic guiding neighboring proteins to cooperatively occupy chromatin fibers. Furthermore, DAF-seq enables the synchronous identification of single-molecule chromatin and genetic architectures – resolving the functional impact of rare somatic variants, as well as transitional chromatin states guiding haplotype-selective promoter actuation. Finally, we demonstrate that single-cell DAF-seq enables the accurate reconstruction of the diploid genome and epigenome from a single cell, revealing that a cell’s accessible regulatory landscape can diverge by as much as 63% while still retaining the cell’s identity. Overall, DAF-seq enables the comprehensive characterization of protein occupancy and chromatin accessibility across entire chromosomes with single-nucleotide, single-molecule, single-haplotype, and single-cell precision.

## Main Text

Gene regulation occurs at the level of an individual chromatin fiber within a single human diploid cell. However, these regulatory patterns can markedly diverge between individual chromatin fibers based on the presence of underlying genetic variants (germline or somatic) or via the coordinated co-occupancy or co-actuation of chromatin features across loci or entire chromosomes. Consequently, resolving the functional output of a chromatin fiber requires the precise mapping of both the genetic composition and protein occupancy of that fiber with single-nucleotide precision. Furthermore, the functional output of each fiber can only be appropriately interpreted within the broader context of how that fiber is structured across a population of cells or relative to the rest of the diploid genome within a single cell.

Although numerous methods exist for resolving single-molecule protein occupancy or single-cell chromatin accessibility, these existing approaches are limited in either their ability to achieve high-resolution protein occupancy patterns^1^, deep targeted enrichment^2,3^, or high-resolution patterns across a single-cell^4,5^. For example, long-read methyltransferase stenciling approaches typically require the direct sequencing of modified DNA, as these methylation marks are not preserved during DNA amplification. Consequently, these approaches typically require millions of cells per reaction and are often limited to providing only 10-1000-fold enrichment. Furthermore, single-cell chromatin assays result in sparse single-cell chromatin data owing to inherent sampling and signal-to-noise limitations with cleavage-based chromatin mapping.

To overcome these limitations, we have developed a single-molecule chromatin fiber sequencing method that is compatible with DNA amplification and provides near-nucleotide resolution (**Fig. 1a**). Specifically, this method leverages a variant of the double-stranded cytidine deaminase toxin A (DddA)^6,7^ to stencil protein occupancy in the form of deaminated cytidines. Upon DNA amplification, these deaminated cytidines create distinct modifications to the DNA sequence, and comparison of these modifications between individual DNA molecules can be used to resolve the genetic composition and protein occupancy of each multi-kilobase chromatin fiber.

**Figure 1.**
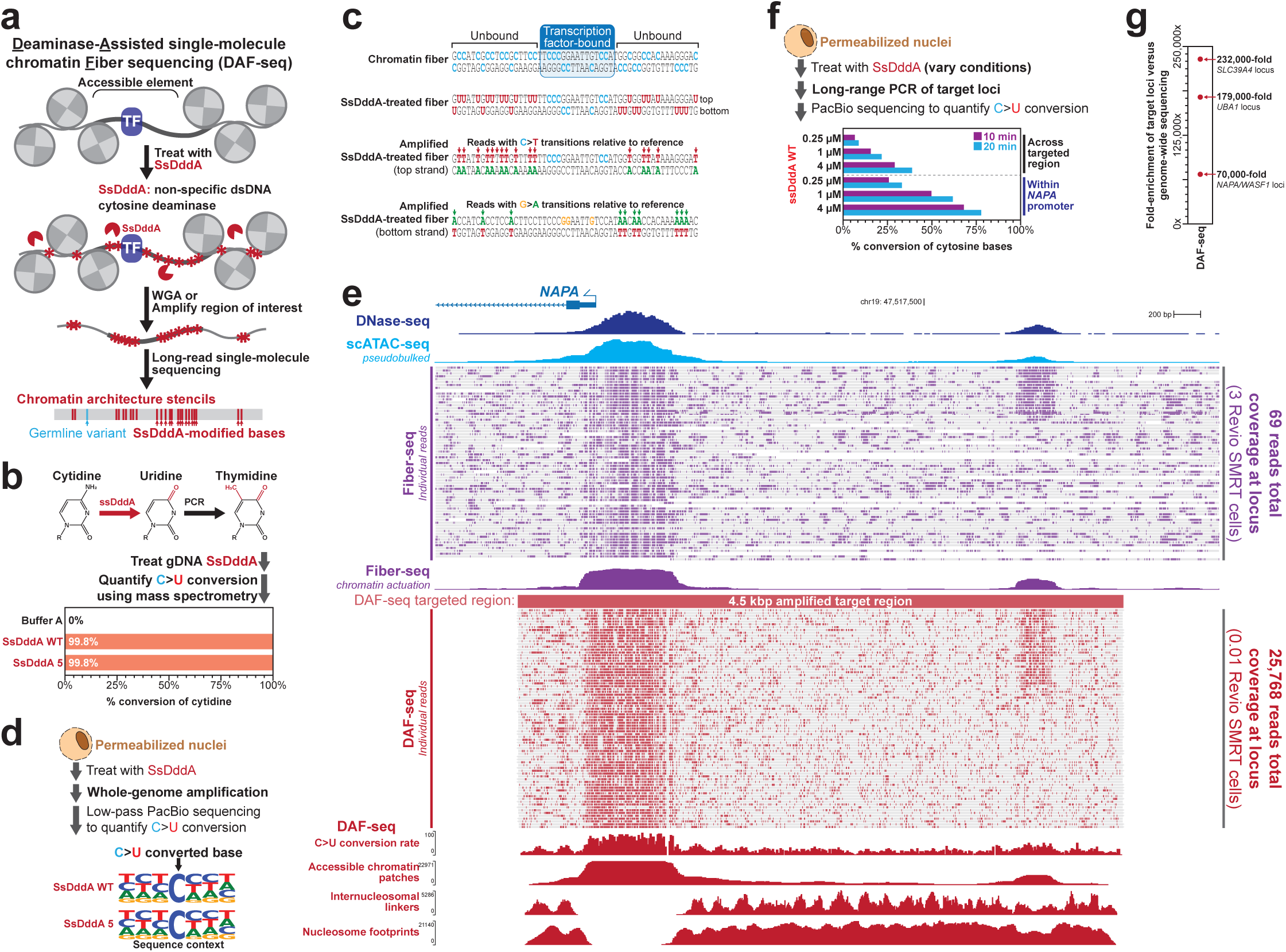
Deaminase-assisted single-molecule chromatin fiber sequencing (DAF-seq). **(a)** Schematic for DAF-seq using the non-specific cytidine deaminase SsDddA to selectively stencil single-molecule protein occupancy using deaminated cytidines. **(b)** Diagram showing deamination of cytidine to uridine and subsequent conversion to thymidine upon PCR amplification. Mass spectrometry quantification of cytidine in untreated genomic DNA, as well as genomic DNA treated with wild-type SsDddA and the variant SsDddA 5. **(c)** Demonstration of how protein occupancy impacts cytidine deamination and how this results in C->T or G->A transitions relative to the reference depending on whether the top or bottom strand is used as the template. **(d)** Motif logos showing the sequence context of cytidine deamination genome-wide after treatment of nuclei with wild-type SsDddA and the variant SsDddA 5. **(e)** Genomic locus showing GM12878 DNase-seq, pseudo-bulked scATAC-seq, Fiber-seq and targeted DAF-seq data at the *NAPA* promoter. Region targeted for amplification noted in red, as well as deamination rate, and coverage of chromatin features derived from the DAF-seq data. Note that DAF-seq used only 1% of a SMRT cell for this sequencing, and only 78/25,768 DAF-seq molecules from this locus are displayed. **(f)** Deamination rate of the targeted NAPA region and promoter element in particular after treatment of GM12878 with various DAF-seq reaction conditions. **(g)** Enrichment of the targeted region versus genome-wide sequencing after targeted DAF-seq of four separate loci in GM12878 cells.

### *Simiaoa sunii* DddA (SsDddA) efficiently and specifically modifies accessible cytidines

Cytidine deaminases with activity towards double-stranded (dsDNA), such as the recently described enzyme DddA, provide a potential solution for mapping chromatin architectures at C·G base pairs. Specifically, SsDddA modifies cytidine to uridine, resulting in C->T mutations upon amplification (**Fig. 1b**), which are readily observable by both short-and long-read DNA sequencing. However, for cytidine deaminases to be optimal for chromatin stenciling, they must have minimal sequence biases, be highly catalytically active, and be highly specific for accessible DNA. Although the originally described DddA enzymes have substantial TC sequence biases^6,8^, subsequent studies employing directed evolution or phylogenetic approaches have identified DddA variants with reduced sequence bias when tethered to a Cas9 enzyme^9–11^. To test whether these enzymes were suitable for chromatin stenciling, we optimized the production of two recombinant DddA variants from the bacterial species *Simiaoa sunii* (SsDddA) using a bacterial system (**Extended Data Fig. 1**) and quantified their cytidine deamination activity on purified double-stranded genomic DNA (gDNA) using mass spectrometry (**Fig. 1b**). This revealed that recombinant SsDddA^7^ is highly catalytically active, deaminating 99.8% of cytidines within gDNA. To confirm that SsDddA has minimal sequence bias within a chromatin context, we treated nuclei with SsDddA (**Fig. 1c**) and performed long-read sequencing of whole-genome amplified DNA from these SsDddA-treated nuclei, similarly showing cytidine deamination with no sequence bias (**Fig. 1d**). Furthermore, we quantified the impact of 5-methylcytosine (5mC) on SsDddA activity using DNA templates treated with the CpG methyltransferase M.SssI, demonstrating that SsDddA can deaminate 5mCpG, albeit with reduced activity (**Extended Data Fig. 2**).

To determine the optimal SsDddA reaction conditions for chromatin stenciling, we treated GM12878 nuclei with a range of enzyme concentrations and treatment times and evaluated each condition using long-read sequencing of targeted amplicons from two genomic loci, the *NAPA* and *WASF1* promoters, that have extensive paired DNase-seq and single-molecule chromatin fiber sequencing (Fiber-seq) data (**Fig. 1e**). Using 1/100th of a Revio SMRT cell we sequenced these regions to 25,672x coverage for *NAPA* and 46,264x for *WASF1*. Cytidine deamination was highly specific to orthogonally defined regulatory elements and internucleosomal linker regions in a manner that mirrored the paired DNase-seq and Fiber-seq data (**Fig. 1e**). At the highest enzyme concentration (4 μM) we observed >50% deamination within accessible regulatory elements (**Fig. 1f**, **Extended Data Fig. 3**), while still preserving adjacent nucleosome footprinting patterns – enabling the near nucleotide precise mapping of protein occupancy events, especially in cytidine rich CpG islands (**Fig. 1e**). Furthermore, we observed that each molecule’s deamination pattern could be readily used as a unique molecular identifier (UMI) to identify reads arising from PCR duplicates (**Extended Data Fig. 4**) due to a practically countless combination of C->T mutations that could exist along a 5 kb fiber. Furthermore, application of targeted DAF-seq to two additional loci resulted in a 70,000 to 230,000-fold enrichment relative to untargeted genome-wide chromatin stenciling approaches (**Fig. 1g**), with the majority of sequenced reads constituting unique molecules at a sequencing depth of 100,000 (**Extended Data Fig. 4**). Overall, these findings demonstrate that recombinant SsDddA can be readily purified using a bacterial system, has minimal sequence bias, is highly catalytically active, and is highly specific for accessible DNA—features that make it well suited for chromatin stenciling with single-molecule and single-nucleotide precision.

### DAF-seq disentangles the regulatory logic of single-molecule TF co-occupancy

The single-molecule and single-nucleotide precision of DAF-seq combined with its high-sequencing depth offers the potential to precisely delineate how individual TFs occupy and co-occupy a given regulatory element. To explore this, we identified 11 high-confidence TF binding elements within the *NAPA* promoter and quantified their per-molecule occupancy (**Fig. 2a,b**), demonstrating that the occupancy of each element ranged from 13% (element 6) to 96% (element 11) of fibers (**Fig. 2c**). We observed that single-fiber co-occupancy for each pair of binding elements is relatively homogeneous, with most co-occupancy deriving from the highly bound element 11 (96% of fibers), which contains a CTCF binding element (**Fig. 2d**). We next determined whether the occupancy of any of these binding elements was codependent, which can be accomplished by calculating the difference between the proportion of fibers co-bound at a pair of elements (*i.e.,* ‘observed co-occupancy’) and the product of the separately bound propositions at each of these two elements (*i.e.,* ‘expected co-occupancy’). Elements with an observed co-occupancy significantly greater than its expected co-occupancy can suggest potential cooperative co-binding activity. Using this, we observed highly codependent occupancy between elements 1, 2 and 3 (**Fig. 2e**).

**Figure 2.**
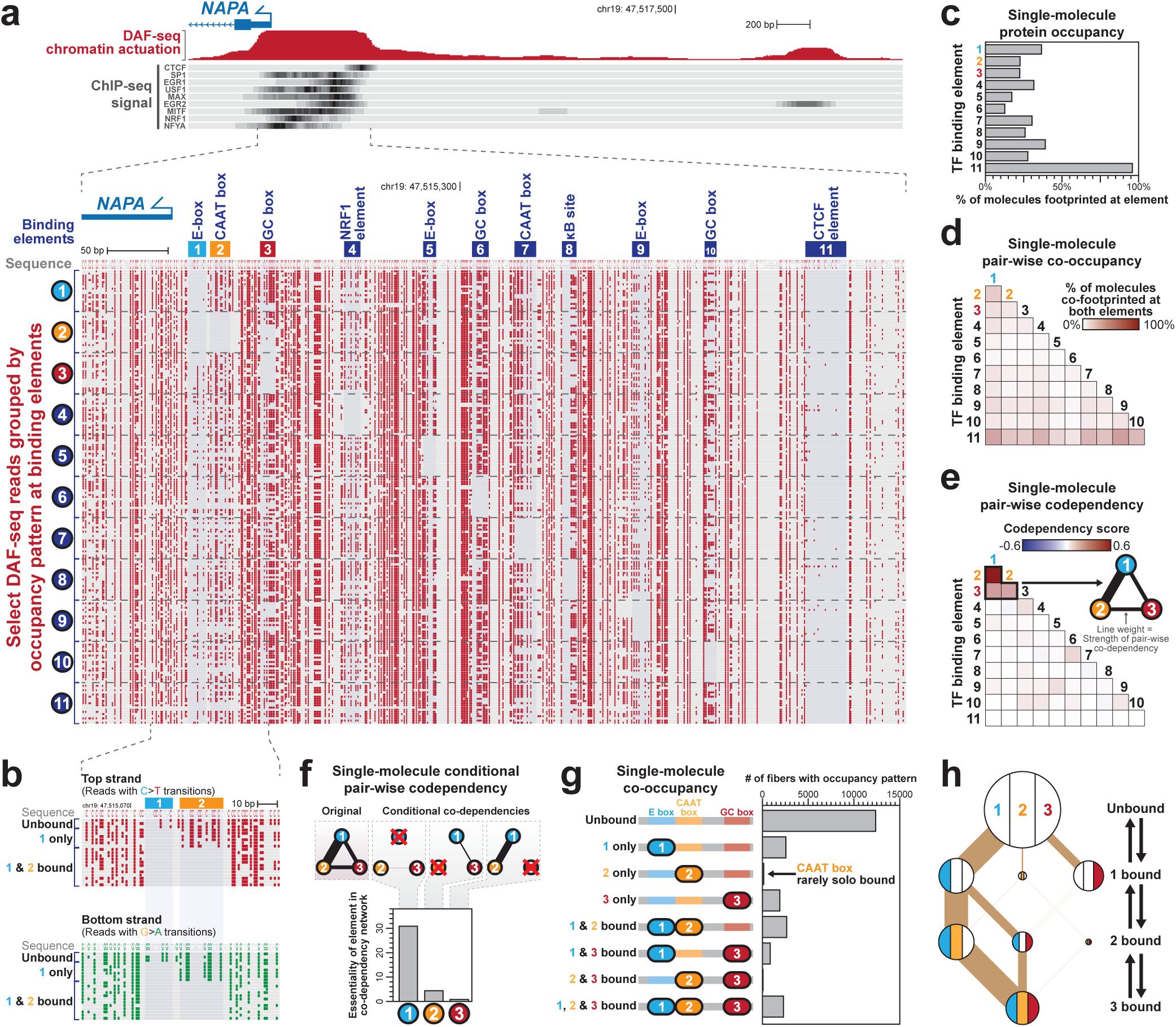
DAF-seq disentangles the regulatory logic of single-molecule TF co-occupancy. **(a)** (top) Targeted DAF-seq chromatin actuation and ChIP-seq data from the *NAPA* promoter in GM12878 cells. (bottom) Zoom-in of the *NAPA* promoter showing single-molecule DAF-seq profiles with deaminated bases marked in red. Only a subset of ‘top-strand’ reads are shown, with reads clustered based on their occupancy pattern at 11 well-defined TF binding elements within the *NAPA* promoter. **(b)** Zoom-in of DAF-seq data at elements 1 and 2 with both ‘top-strand’ and ‘bottom-strand’ reads shown. **(c)** Bar graph showing single-molecule protein occupancy measurements of the 11 binding elements within the *NAPA* promoter. **(d)** Heatmap showing single-molecule pair-wise TF co-occupancy of the 11 elements within the *NAPA* promoter. **(e)** Heatmap showing single-molecule pair-wise TF codependency of the 11 elements within the *NAPA* promoter, as well as a network diagram of elements 1, 2, and 3 with the edge weights corresponding to the strength of the codependency. **(f)** Single-molecule conditional codependency with a bar graph showing the impact that removal of occupancy at elements 1, 2, or 3 has on the codependency score of the remaining elements within this cluster. Higher score indicates that an element is essential for a codependent network. **(g)** Bar graph showing the number of molecules with occupancy at different combinations of elements 1, 2, or 3. **(h)** Directed network diagram showing how elements 1, 2, or 3 go from the unbound to fully bound state. Individual nodes are colored based on their occupancy patterns and are sized based on the data from panel g. The edges connecting the unbound and fully bound state are weighted based on the size of the smallest node through which they traverse during this path, and the translucency of each edge leaving the 1 bound state is dependent upon the data from panel f.

To discern which elements potentially drive these interactions, we employed a network graph approach to quantify the conditional codependency between these three elements^12^. We quantified the change in the graph’s overall codependency when selectively analyzing only fibers unbound at a given element, resulting in scores representative of the essentiality of each element for codependent TF occupancy within the graph (**Fig. 2f**). Element 1 was by far the most essential element, supporting a clear directional effect whereby element 1 drives the codependent occupancy of these three elements. In addition, whereas element 1 can be bound on its own, >96% of fibers bound at element 2 are concurrently bound at element 1, further supporting a directional codependency between these elements (**Fig. 2g**). These directional graphs enabled us to implicate element 1 in nucleating co-occupancy across elements 1, 2, and 3 within the *NAPA* promoter *in vivo* (**Fig. 2h**). Of note, whereas element 1 contains a predicted high-affinity element for USF1 & USF2, the CAAT box in element 2 contains a predicted low-affinity sequence element for NFY-A, consistent with distance-dependent cooperative binding between USF1/2 and NFY-A, whereby USF1/2 binds and recruits NFY-A through its USF Specific Region (USR) domain^13–15^. Overall, these findings demonstrate marked heterogeneity within the single-molecule occupancy of TFs within the *NAPA* promoter, with the majority of these TFs (8/11) binding in a largely independent manner.

### Synchronous single-molecule genomic and chromatin profiles

We next evaluated the ability of DAF-seq to disentangle SsDddA-induced deaminations from germline genetic variants, leveraging the fact that SsDddA only modifies one strand of a C/G base-pair, and that reads can be readily partitioned into ‘top-strand’ and ‘bottom-strand’ reads based on their predominance of C->T versus G->A changes, respectively, relative to the reference (**Fig. 1c**). Specifically, at a C/G base pair, although the top strand may be variably deaminated and converted to a T, the bottom strand will always remain a G, enabling one to readily distinguish whether a position was originally a C/G or T/A base-pair (**Fig. 3a**). Consequently, DAF-seq reads mapping to a reference position that has a C/G base-pair on both haplotypes will contain 100% G on the bottom-strand, and between 0 and 100% C on the top strand, with the base content of the top strand reflecting the frequency by which that site is deaminated (**Fig. 3a**). Overall, this approach enabled the accurate delineation of germline variants, as well as the haplotype phasing of DAF-seq reads based on these germline variants. The one exception comes when the only heterozygous germline variant in a read is a C/T or G/A variant, allowing DAF-seq reads from only one of the two strands to be accurately phased (*i.e.,* the bottom or top strand, respectively). To validate this, we applied targeted DAF-seq to a 4.4 kb region containing only a single C/T heterozygous variant (rs56269549) in GM12878 cells, allowing us to successfully phase bottom strand DAF-seq reads. This targeted region is located on the X chromosome and spans four *UBA1* transcriptional start sites (TSSs), three of which are selectively accessible on only one of the haplotypes (*i.e.,* Xa) in a population of GM21878 cells with allelically skewed X chromosome inactivation^16^. This haplotype-resolved DAF-seq data demonstrated that whereas the canonical *UBA1* TSS is accessible on both the Xa and Xi, the three upstream TSSs are selectively accessible on only the Xa, confirming the ability of DAF-seq to accurately haplotype phase reads, even when they contain only a single heterozygous C/T variant (**Fig. 3b,c**).

**Figure 3.**
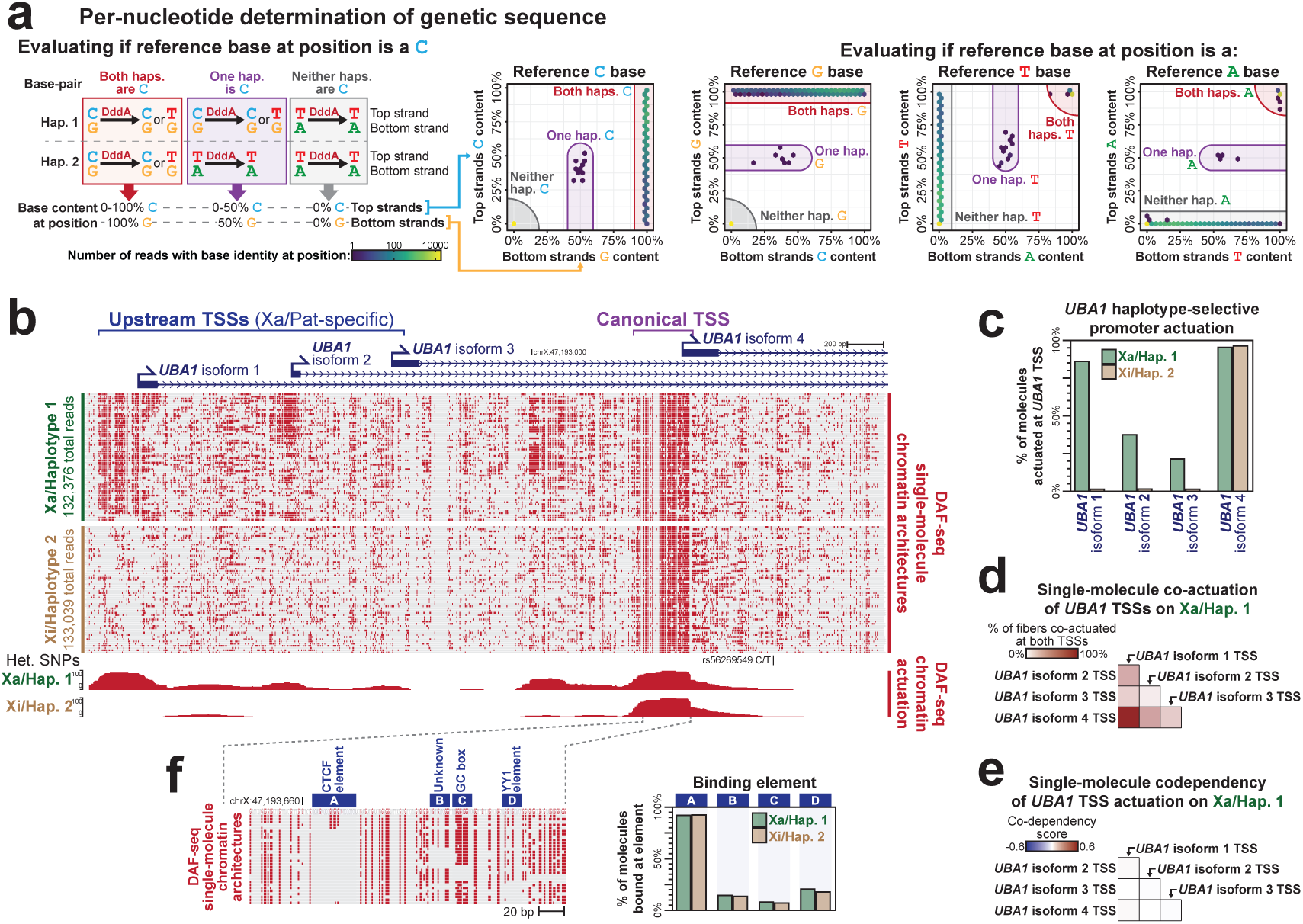
Simultaneous single-molecule genomic and chromatin profiles. **(a)** (left) Diagram showing the evaluation of the underlying genetic architecture at each reference genome position. Specifically, reads are divided into ‘top-strand’ and ‘bottom-strand’ based on their predominance of C->T versus G->A mutations relative to reference. Below, the base content at a position at all reads from the top and bottom strands are quantified and used to evaluate the germline genetic content at that position. (right) Hexbin plots showing the base content at all of the targeted DAF-seq regions used in this manuscript. **(b)** Haplotype-phased single-molecule and aggregate targeted DAF-seq data of the *UBA1* promoter in GM12878 cells which have allelically skewed XCI. The single C/T germline variant used for phasing is indicated. **(c)** Bar graph showing chromatin actuation at each of the four *UBA1* TSSs by haplotype. **(d,e)** Heatmap showing single-molecule pair-wise co-actuation (d) and codependency (e) of the four *UBA1* TSSs along the Xa. **(f)** (left) Zoom-in of *UBA1* isoform 4 TSS showing single-molecule protein occupancy at four binding elements. (right) Bar graph showing single-molecule protein occupancy at each of the four *UBA1* isoform 4 TSS binding elements by haplotype.

Leveraging this haplotype-phased data, we next sought to determine if these *UBA1* TSSs are independently actuated or if their actuation shows codependency. Importantly, we limited this analysis to only reads arising from the Xa, enabling us to determine if any observed codependency in the unphased data is simply a reflection of X inactivation (**Extended Data Fig. 5**), or potentially reflects cooperative behaviors between the various TSSs on a single-molecule level. Although these four *UBA1* TSSs showed strong co-actuation along the Xa (**Fig. 3d**), the actuation of each of these TSSs was largely occurring independently of each other (**Fig. 3e**). Overall, this indicates that the Xa chromatin environment is suitable for the actuation of *UBA1* TSSs and that the actuation of one of these TSSs neither positively nor negatively influences the ability of the other three TSSs to become actuated along that fiber.

These findings demonstrated a <1 kb boundary separating chromatin that is subjected to X chromosome inactivation (XCI) (*i.e.,* the *UBA1* isoform 3 TSS) versus XCI escaping (*i.e.,* the *UBA1* isoform 4 TSS). To determine if the ability of the *UBA1* isoform 4 TSS to escape XCI is mediated by differential TF usage between the Xa and Xi we compared TF occupancy patterns at four binding elements within this TSS (**Fig. 3f**). Notably, we observed only subtle differences in TF occupancy patterns between the Xa and Xi, indicating that TF occupancy patterns within the *UBA1* isoform 4 promoter are not establishing the ability of the upstream region to differentially act as a boundary for XCI on the Xa and Xi.

### *SLC39A4* haplotypes modulate single-molecule promoter actuation patterns

Over 4 million germline genetic variants showing a significant association with gene expression patterns in at least one tissue have been identified^17^. However, resolving the mechanisms that mediate these associations is quite challenging, especially for tissue-specific associations with a relatively low effect size. For example, the rs2280838-T haplotype is associated with modestly increased *SLC39A4* transcript levels in liver tissue (**Fig. 4a**), but the mechanisms driving this are unknown. Importantly, promoter variants in *SLC39A4* have recently been shown to cause acrodermatitis enteropathica (MIM: 201100)^18^, so understanding how common haplotypes modulate the expression of this gene can have implications for designing therapeutics for this condition. To address this, we applied targeted DAF-seq to a 2.1 kb region spanning the *SLC39A4* promoter from two samples heterozygous at rs2280838(C/T): primary human liver tissue, which expresses *SLC39A4*, and GM12878 lymphoblastoid cells, which do not express *SLC39A4* (**Fig. 4a**). We sequenced to a depth of ∼1,200,000x targeted coverage in each of these samples, combined the DAF-seq reads from both tissues, and clustered them by their single-molecule deamination patterns (**Fig. 4b**, **Extended Data Fig. 6**). This exposed distinct patterns between clustered reads in their nucleosome positioning patterns and actuation status of the *SLC39A4* promoter (**Fig. 4c**). Notably, the contribution of reads from the two tissues and haplotypes markedly differed between the clusters (**Fig. 4c**). Specifically, clusters 4, 5, and 6, which consist of variably positioned nucleosome arrays, were selective to reads from lymphoblastoid cells. In contrast, clusters 2 and 3, which consist of nucleosome arrays with exquisitely well-positioned nucleosomes within the *SLC39A4* promoter, were represented by reads from both liver and lymphoblastoid cells, with a preference for liver reads from the rs2280838-C haplotype. Notably, cluster 1 alone showed actuation of the *SLC39A4* promoter, and consisted almost exclusively of chromatin fibers from liver tissue, with 72% of those reads originating from the rs2280838-T haplotype (**Fig. 4c**), indicating that the association of rs2280838 with *SLC39A4* transcript levels in liver is likely mediated through differences between the two haplotypes in their propensity for forming actuated chromatin at the *SLC39A4* promoter.

**Figure 4.**
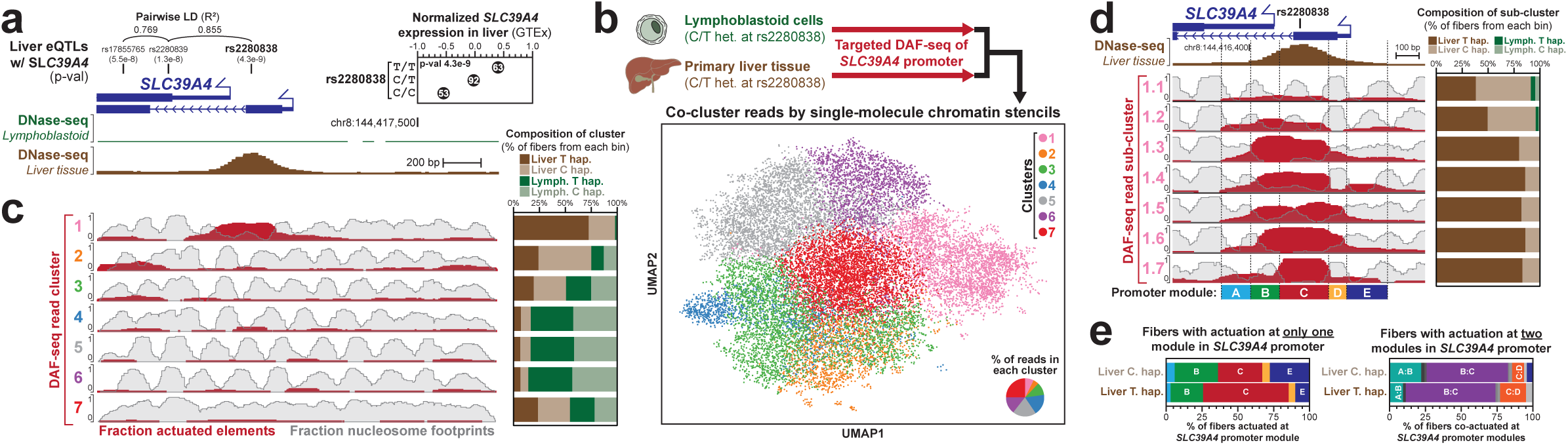
*SLC39A4* haplotypes modulate single-molecule promoter actuation patterns. **(a)** Liver eQTL data from GTEx of three single nucleotide polymorphisms (SNPs) within the *SLC39A4* promoter as well as their linkage disequilibrium using 1,000 genomes data. Below are DNase-seq data at this region in GM12878 cells and primary liver tissue. **(b)** (top) Diagram showing targeted DAF-seq on the *SLC39A4* promoter in two cell types heterozygous for rs2280838 followed by the Leiden clustering of reads based on their deamination profiles and subsequent projection of these reads into UMAP space. (bottom-right) Pie chart showing the relative contribution of each cluster to the total number of reads. **(c)** (left) Aggregate chromatin actuation and nucleosome occupancy of reads from the different clusters identified in panel b. (right) Stacked bar chart showing the relative contribution of liver or GM12878 reads to the different clusters, as well as the relative contribution of rs2280838-C or rs2280838-T. **(d)** Results from sub-clustering of the reads from cluster 1 in panel b. (left) Aggregate chromatin actuation and nucleosome occupancy of reads from the different subclusters. (right) Stacked bar chart showing the relative contribution of liver or GM12878 reads to the different sub-clusters, as well as the relative contribution of rs2280838-C or rs2280838-T. (bottom) Promoter modules defined using above aggregate profiles. **(e)** Identity of actuated module split by haplotype for fibers with chromatin actuation at only one of the 5 modules. **(f)** Similar to panel e, but for fibers with actuation at only 2 of the 5 modules.

To explore how rs2280838 modulates *SLC39A4* promoter actuation, we isolated the fibers from cluster 1 and re-clustered them (**Extended Data Fig. 6**), revealing distinct single-molecule patterns of focal chromatin actuation within the *SLC39A4* promoter that are hidden from short-read based chromatin assays (**Fig. 4d**). Specifically, the *SLC39A4* promoter appeared to be naturally subdivided into five distinct modules that were variably actuated across the individual fibers. Notably, the module containing rs2280838, module C, corresponds to the same position occupied by an exquisitely well-positioned nucleosome within the *SLC39A4* promoter along non-actuated reads from the liver, raising the prospect that rs2280838 may modulate the occupancy of this nucleosome during *SLC39A4* promoter actuation. To further delineate how the chromatin architecture at the rs2280838 variant differs by allele, we quantified the proportion of fibers with 0, 1, 2, 3, 4, or 5 modules actuated (**Extended Data Fig. 6**). Notably, rs2280838-C had a higher proportion of fibers with 0 modules actuated compared to rs2280838-T, corresponding to a closed chromatin state which is consistent with reduced expression from the rs2280838-C allele. In contrast, rs2280838-T was enriched for 3 actuated modules, suggesting that the 3-module actuation state is conducive for transcriptional activation of the promoter. Next, we hypothesized that the compositions of the 1-, 2-, and 3-actuated module states could reflect the mechanism by which individual chromatin fibers transition between the 0-and 3-module actuated states. For the 1-module actuated state, module C was preferentially actuated in rs2280838-T fibers, whereas modules B, C, or E were similarly actuated in rs2280838-C fibers (**Fig. 4e**), consistent with a role for rs2280838 in modulating nucleosome occupancy of module C (**Extended Data Fig. 6**). In contrast, the specific combinations of modules actuated in the 2-and 3-module bound states were more similar between haplotypes and largely included actuation of module C, suggesting a critical role of module C actuation in the efficient transition from a 0-module to 3-module actuated chromatin state. Overall, by simultaneously capturing both the chromatin and genomic profile of individual chromatin fibers, we reveal a mechanism by which accessibility and transcriptional activity of the *SLC39A4* TSS is potentiated by module C actuation along the rs2280838-T haplotype.

### Resolving the functional impact of rare non-coding mosaic mutations

Our understanding of the prevalence and functional impacts of somatic variation is constrained by inherent limitations with short-read chromatin assays in co-measuring both genetic and chromatin information at targeted loci. We have recently demonstrated that single-molecule chromatin assays are well suited for measuring the functional impacts of mosaic variants with a high variant allele fraction (VAF)^19^, and given the ability of targeted DAF-seq to synchronously measure single-molecule genomic and chromatin profiles at extremely high depth and with high accuracy, we hypothesized that this approach could enable the interrogation of the functional impacts of low VAF somatic variants.

To test this, we leveraged a cell-based model of somatic variation, which includes a 49:1 mixture of the B lymphoblast cell line COLO829BL (BL), and a melanoma tumor line COLO829T (T) derived from the same individual (**Fig. 5a**). Fiber-seq data from these two unmixed cell lines^16^ identified a CC>TT somatic dinucleotide mutation on one haplotype of COLO829T (chr17:19,447,245-19,447,246) that ablates an overlying CTCF binding element, causing selective loss of CTCF occupancy and chromatin accessibility on the variant haplotype relative to reference (**Fig. 5b**). Application of targeted DAF-seq to a 3.8 kb region spanning the variant in the 49:1 BL:T mixture readily exposed the presence of these somatic variants (**Extended Data Fig. 7**), and identified 1,701 of 115,991 bottom strand (G-to-A) reads as containing the CC>TT variant, for a VAF of 1.5% (**Fig. 5a**). We benchmarked the genetic accuracy of DAF-seq against deep, Illumina PCR-free whole-genome sequencing of the same 49:1 BL:T mixture, which identified 6 of 437 reads with the CC>TT variant for a variant allele fraction (VAF) of 1.4%. However, in addition to accurately quantifying the VAF for this variant, DAF-seq also readily exposed that the variant reads lost chromatin accessibility and nucleosome phasing at the targeted element (**Fig. 5c**). These results showcase DAF-seq as a unique tool for the simultaneous identification and functional characterization of low-frequency somatic variants within human tissues, and demonstrate that nucleosome phasing surrounding a CTCF element is abrogated upon loss of CTCF occupancy and chromatin accessibility.

**Figure 5.**
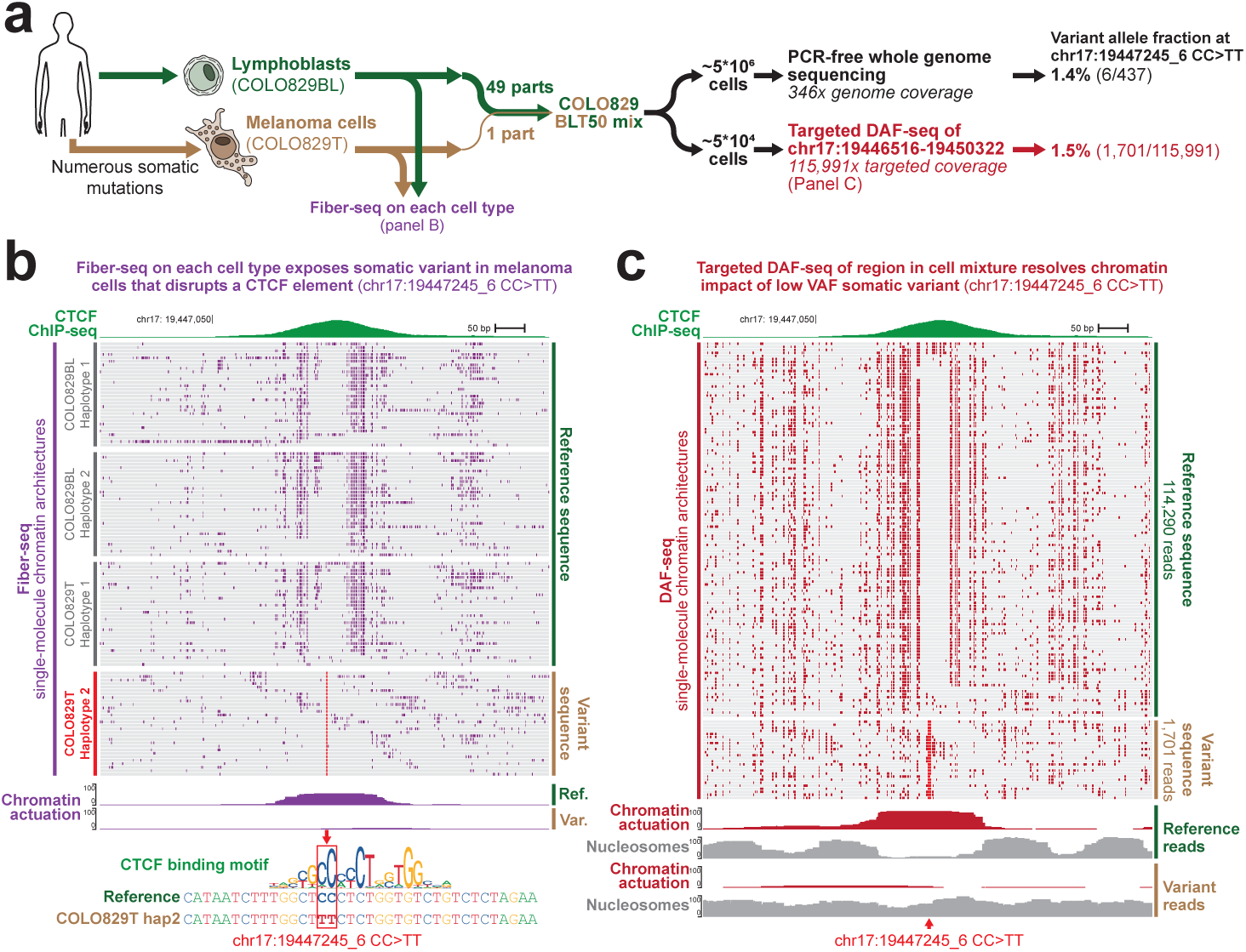
Resolving the functional impact of low VAF mosaic mutations. **(a)** (left) Schematic showing the generation of the COLO829 BLT50 cell mixture, which is a mixture of the lymphoblastoid cell line COLO829BL and the melanoma cell line COLO829T derived from the same individual. Fiber-seq was performed on each of these cell lines separately, as is shown in panel b. (right) Whole genome PCR-free Illumina sequencing and targeted DAF-seq of the COLO829 BLT50 mixture showing the VAF at the chr17:19447245_6 CC>TT variant. **(b)** Fiber-seq data from the COLO829BL and COLO829T cells showing the per-molecule and aggregate chromatin actuation data surrounding the chr17:19447245_6 CC>TT variant, as well as the impact of this variant on a CTCF binding element. Note the complete loss of per-molecule CTCF occupancy and chromatin actuation on the reads containing the variant sequence. **(c)** Targeted DAF-seq of the same region as panel b in the COLO829 BLT50 cell mixture showing the single-molecule and aggregate chromatin patterns on reads containing the reference sequence (top) and the chr17:19447245_6 CC>TT variant (bottom). Note the complete loss of chromatin actuation and nucleosome positioning along the variant reads.

### Ultra-long consensus reads and chromosome-scale genomic phasing in single cells

One of the major drawbacks of targeted DAF-seq is that chromatin fiber lengths are limited by long-range PCR. However, the regulatory elements modulating a gene can be located >1 Mb away^20^, which is well beyond the ability of long-range PCR or long-read PacBio or Oxford Nanopore Technologies (ONT) sequencing to routinely capture. Single-cell DAF-seq (scDAF-seq) has the potential to overcome this, as unlike enzymatic methylation^1–3^, DAF-seq chromatin stencils are preserved during amplification, enabling the generation of a sufficient amount of DNA from a single cell for long-read sequencing. To evaluate DAF-seq as a single-cell sequencing technology, we treated permeabilized GM24385 (HG002) cells with SsDddA and sorted cells into individual sample wells using fluorescence-activated cell sorting (FACS). We then performed whole-genome amplification (WGA) separately on each cell using primary template-directed amplification (PTA)^21^ and sequenced the amplification products from four cells with PacBio Hifi long-read sequencing (**Fig. 6a**). Three cells were sequenced to a depth of 22.1-29.1 Gb, while the fourth cell was more deeply sequenced to 78.3 Gb (N50 for sequencing fragment length of 4.2-5.4 kb) (**Fig. 6b,c**). Critically, PTA’s increased preference for priming on the primary template produced numerous partially overlapping amplicons originating from the same template, with all autosomes containing four ‘haplotype-strand’ templates at each genomic position, corresponding to the top and bottom strands of each of the two haplotypes (**Extended Data Fig. 8**). Each ‘haplotype-strand’ template has a unique deamination pattern that reflects the strand-specific occupancy pattern of proteins at that site within that cell. As the precise locations of deaminase-induced mutations are stochastic due to incomplete SsDddA deamination and heterogeneous nucleosome positioning, we reasoned that we could generate consensus reads along each ‘haplotype-strand’ that were comprised of individual sequencing reads that uniquely overlapped based on their deamination patterns (**Fig. 6a**), analogous to generating consensus reads from multiple sequencing passes during circular consensus sequencing (CCS).

**Figure 6.**
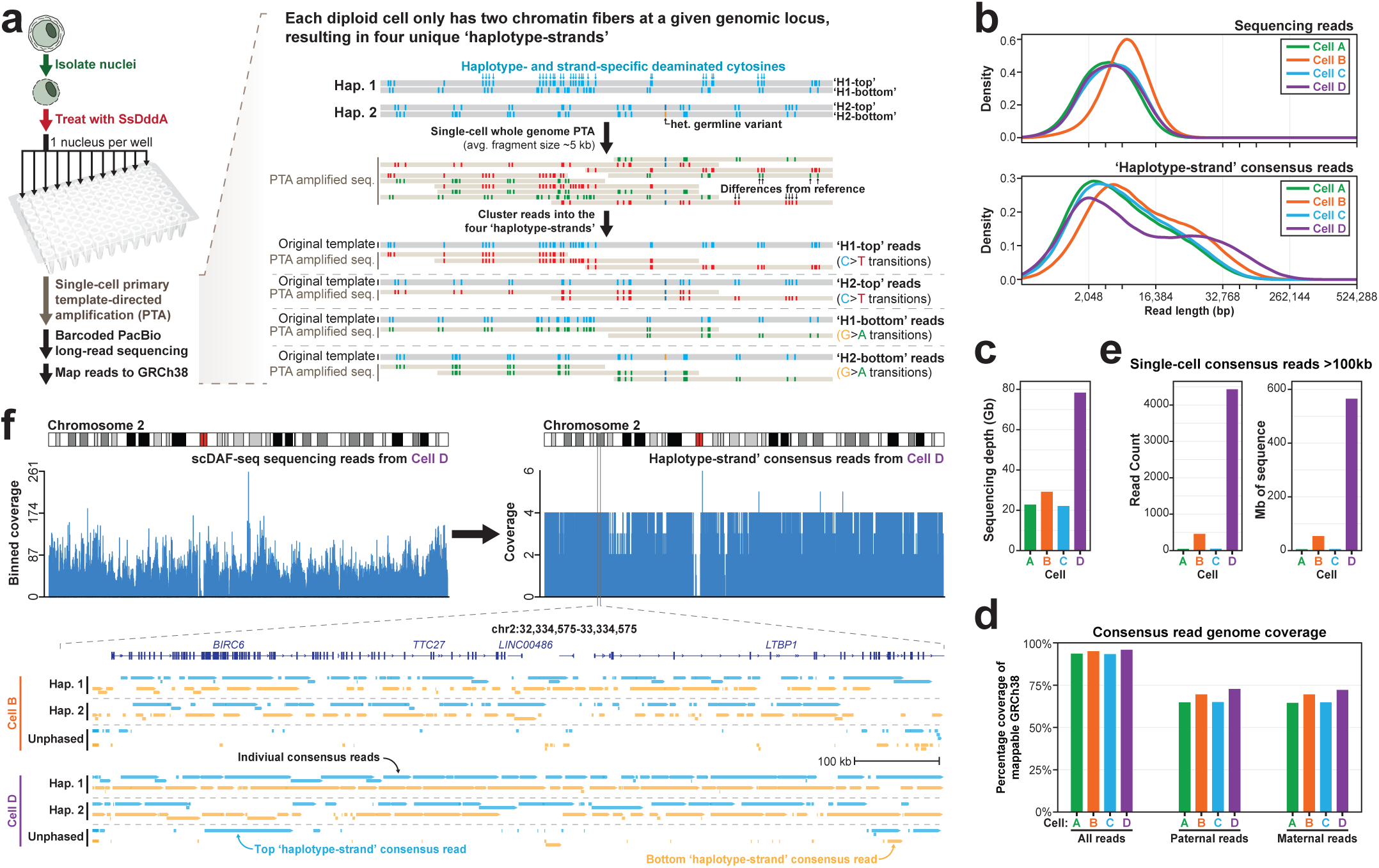
Chromosome-scale genomic phasing in single cells. **(a)** Schematic for single-cell DAF-seq. Specifically, permeabilized cells are treated with SsDddA and then sorted into individual wells of a plate, with each well being subjected to PTA. PTA reads from each well are then sequenced using PacBio HiFi sequencing and mapped to GRCh38. Individual reads are then identified as arising from either the ‘top’ or ‘bottom’ strand based on their pattern of either C>T or G>A mutations relative to the reference, respectively. Overlapping reads are then combined to generate consensus reads for each ‘haplotype-strand’, which are then haplotype-phased across the entire genome using parental short-read data. **(b)** (top) Density plot showing the size distribution of sequencing reads (top) or ‘haplotype-strand’ consensus reads (bottom) from the four cells subjected to scDAF-seq. **(c)** Sequencing depth of each cell in terms of gigabases (Gb). Note that only a fraction of the library was sequenced for each cell. **(d)** Genomic coverage of consensus reads for all reads, as well as haplotype-phased reads. **(e)** Number of consensus reads >100,000 bp in length, as well as the amount of genomic sequence covered by these ultra-long reads in each cell. **(f)** (top) Genomic coverage of chromosome 2 from the raw sequencing reads from cell D, as well as the coverage from the consensus reads from this same cell. (bottom) Genomic locus showing the consensus reads from cell B and D, with reads colored based on whether they are from the top or bottom strand, and split based on their phasing.

To accomplish this, we mapped reads from each cell to GRCh38 and collapsed overlapping reads originating from the same ‘haplotype-strand’ template to generate individual ‘consensus reads’ for each strand and haplotype of that cell (methods). In the cell with the highest sequencing depth, this approach enabled the construction of consensus reads with an N50 of 33.2 kb (Fig. 6b), with at least one read spanning 96% of the mappable, autosomal portions of GRCh38 (2.53 Gb) (**Fig. 6d**). Notably, 4,429 consensus reads from that cell were >100 kb in length, with these single-cell ultra-long consensus reads covering 565 Mb of genomic sequence within that cell (**Fig. 6e**). Furthermore, 64% of all ‘consensus reads’ from the four cells could be accurately phased to their respective haplotype using parental short-read data, resulting in the chromosome-scale haplotype-phasing of 72% of the genome from a single cell (**Fig. 6d**), with a haplotype switch error rate of 2.3-3.3%, a rate comparable to current assembly methods with less than 10-fold coverage^22–24^. Consensus reads demonstrated even coverage across the genome relative to their original underlying sequencing reads (**Fig. 6f**), readily exposing anomalous autosomal genomic loci harboring >4 ‘haplotype-strand’ templates, indicative of possible duplications present in that cell relative to GRCh38. Together, these data demonstrate that scDAF-seq enables the accurate reconstruction of the chromosome-scale haplotype-phased diploid genome from a single cell, with thousands of consensus reads >100,000 bp in length.

### The near-complete chromatin epigenome of a single-cell

Our current understanding of the chromatin epigenome of a single cell is derived from single-cell ATAC-seq, which typically yields only 10,000-20,000 paired-end reads per cell, which correspond to Tn5 insertion sites. This means that the chromatin state of each cell is only resolved at the 9bp surrounding the 20,000-40,000 cut sites (0.01% of the mappable genome)^4,5,25^. In contrast, scDAF-seq enables the genetic evaluation of 96% of each cell’s mappable haploid genome, offering the potential for single-nucleotide precise maps of protein occupancy within a single cell—a ∼7,000-fold improvement in our understanding of a cell’s chromatin architecture. Using the deamination patterns from each consensus read within our scDAF-seq data, we observed that 19-25% of the cytosine bases within each cell were deaminated (**Fig. 7a**), with the predominant pattern of deamination within each cell being driven by nucleosome occupancy (**Fig. 7b**). Notably, unlike the Hia5 m6A-methyltransferase used in Fiber-seq^26^, SsDddA does appear to have some minor enzymatic activity for nucleosome-bound DNA, a property that likely positively contributes to the ability of scDAF-seq to generate consensus reads. In addition to nucleosomes, single-cell deamination patterns clearly demarcated closed and actuated regulatory elements (**Fig. 7c**), as well as individual TF occupancy events within these actuated elements with single-cell and single-nucleotide precision (**Fig. 7d**).

**Figure 7.**
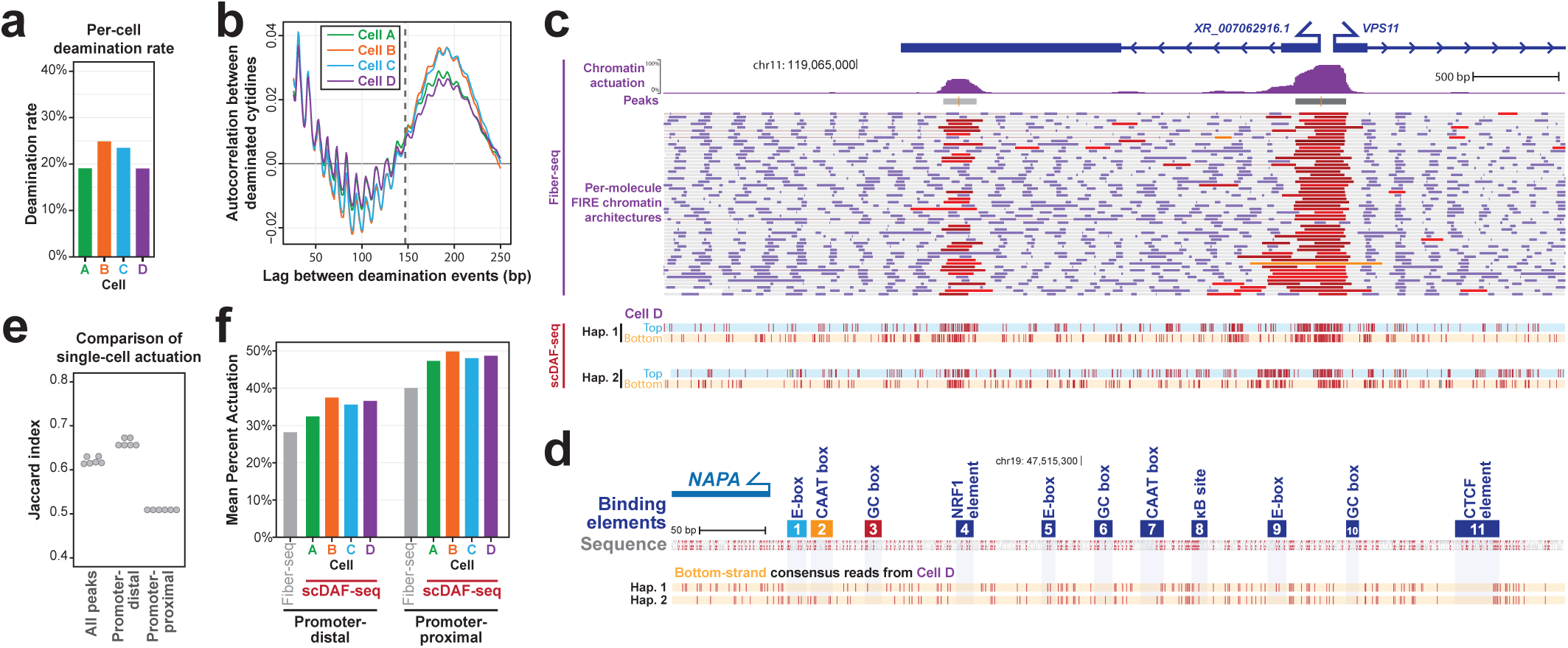
Single-molecule chromatin epigenome of a single-cell. **(a)** Bar graph showing the deamination rate from each of the four sequenced cells. **(b)** Autocorrelation plot of single-molecule deamination patterns in each cell showing a pattern consistent with nucleosomes being the predominant modulator of single-cell and single-molecule deamination by SsDddA. Vertical dashed line is at 147bp. **(c)** Genomic locus showing single-molecule and aggregate Fiber-seq data at the *VPS11* promoter in GM12878 cells (top), as well single-molecule deamination patterns along all four of the ‘haplotype-strand’ consensus reads at this position within Cell D. **(d)** Genomic locus showing single-molecule deamination patterns along the two bottom-strand consensus reads at the same *NAPA* promoter position shown in Figure 2, as well as the position of TF binding elements within this promoter. **(e)** Swarm plot displaying the Jaccard index comparing single-molecule scDAF-seq actuation patterns between cells at genomic loci that are covered across both cells. Elements are defined using paired Fiber-seq data and are further divided into promoter-proximal and promoter-distal peaks. **(f)** Mean single-cell percent actuation of promoter-distal and promoter-proximal elements within each of the four cells subjected to scDAF-seq, as well as from paired bulk Fiber-seq data.

Using this first-ever map of the near complete chromatin epigenome of a human diploid single cell, we sought to evaluate the heterogeneity of chromatin architectures at the single-cell level. Bulk Fiber-seq performed on GM24385 cells identified 154,070 actuated regulatory elements within mappable autosomal regions, 94.9% of which were covered by a consensus read in our deeper sequenced GM24385 cell. Overall, we observed that only 57.6% of these regulatory elements (84,275/146,190) were, in fact, actuated within this single cell, with the specific regulatory elements actuated differing by as much as 63% between the four cells (**Fig. 7e**). Promoter-proximal elements were 36.8% more likely to be actuated at the single-cell level than promoter-distal elements, consistent with their bulk chromatin actuation profiles (**Fig. 7f**). Furthermore, 33% of the elements along haplotype-phased consensus reads demonstrated discordant single-cell actuation patterns (*i.e.,* the element is open on one haplotype and closed on the other within the same cell). Notably, only a minority of these elements represent sites with consistent haplotype-selective chromatin across GM24385 cells, as measured using bulk Fiber-seq (**Extended Data Fig. 9**), demonstrating widespread plasticity in a single-cell’s regulatory landscape.

## Discussion

We present DAF-seq as a transformative method for studying the function of the non-coding genome, providing unprecedented high-resolution targeted single-molecule chromatin fiber sequencing, as well as the first-ever comprehensive maps of the diploid genome and chromatin epigenome from a single cell. We demonstrate that DAF-seq can be performed on frozen primary human tissues as well as cultured cells, and can be used with as little as a single cell as starting material. As DAF-seq uses amplicon-based sequencing, it is compatible with all current short-or long-read sequencing platforms and is well suited for deep profiling of the chromatin architectures of individual regulatory elements—exposing cooperative protein occupancy (**Fig. 2**), chromatin actuation transition states (**Fig. 4**), and the functional impact of low VAF somatic variants (**Fig. 5**).

DAF-seq resolved chromatin transition states can be used to reconstruct how chromatin transitions to and from the closed and open states (**Fig. 4**), as well as the process by which individual TFs co-bind a given regulatory element (**Fig. 2**). As these transition states capture the inherent heterogeneity in chromatin actuation and TF occupancy, they can disentangle the conditional codependency between individual TF binding elements and regulatory elements, offering insights into the structure of regulatory networks that to-date have largely necessitated time-consuming genome or epigenome editing methods to resolve^27,28^. Furthermore, scDAF-seq enables the interrogation of single-molecule regulatory element co-actuation patterns across entire chromosomes, dramatically expanding the ability of single-molecule stenciling to resolve the regulatory logic of gene regulatory networks in diploid organisms.

Targeted DAF-seq and scDAF-seq dramatically improve how we study somatic variants and their functional impact. Specifically, as the deamination pattern on each read can be used as a UMI, PCR duplicates can be readily identified and leveraged to identify a consensus sequence of the primary template, analogous to duplex sequencing methods^29^. Consequently, targeted DAF-seq enables the accurate identification and quantification of low VAF somatic variants (**Fig. 5**) using an economical experimental design (*i.e.,* genomic PCR followed by amplicon sequencing using ∼1% of a PacBio SMRT cell per target). Furthermore, as targeted DAF-seq is compatible with long-read sequencing, it can be leveraged to interrogate somatic variants in complex genomic regions. However, unlike ddPCR^30^ or other amplicon-based approaches, targeted DAF-seq enables the simultaneous quantification of somatic genetic variants, as well as their functional impact on chromatin patterns with improved resolution and throughput relative to existing targeted methods^19^ or genome-wide methods^31,32^.

In addition, scDAF-seq offers the potential to dramatically improve our understanding of somatic variation at the level of single cells. Specifically, by separately labeling each ‘haplotype-strand’ template within a cell, scDAF-seq enables the generation of consensus sequences along each ‘haplotype-strand’ template, thereby correcting for any erroneous sequencing or amplification-induced mutations. This is enabled by the high sequencing accuracy of PacBio HiFi sequencing combined with the high fidelity and primary template preference provided by PTA. The process of generating consensus template sequences dramatically improves the read coverage within a single cell, readily exposing genomic locations with >4 ‘haplotype-strand’ templates that correspond to duplications in that cell relative to the reference. In addition, as each strand is separately interrogated, scDAF-seq offers the potential to readily identify strand-specific mutations at the single-cell level. Finally, we demonstrate that scDAF-seq enables the generation of thousands of ultra-long consensus reads from a single cell, and that the generation of these ultra-long consensus reads can be readily increased by simply sequencing the library from each cell at higher depths. These single-cell ultra-long consensus reads enable the evaluation of single-cell genomic and epigenomic variation within the most complex regions of the genome and lay the groundwork for potentially assembling telomere-to-telomere genomes and chromatin epigenomes from single cells. Overall, DAF-seq dramatically expands our toolkit for single-molecule chromatin stenciling.

## Acknowledgements

We thank Northwest Genome Center and Katherine M. Munson for their assistance in PacBio sequencing, UW Mass Spectrometry Center for their assistance in mass spectrometry experiments, the Fowler lab and Hyeon-Jin Kim for assistance with nuclei sorting, Thomas J. Bell and Kathryn Leonard at The National Disease Research Interchange (NDRI) for primary frozen tissue, and members of the UW-SCRI Somatic Mosaicism across Human Tissues (SMaHT) Genome Characterization Center for generating the COLO829 cell mixture, Illumina sequencing data, and their feedback and support. Funding: A.B.S. holds a Career Award for Medical Scientists from the Burroughs Wellcome Fund and is a Pew Biomedical Scholar. This study was supported by National Institutes of Health (NIH) grants 1DP5OD029630, and UM1DA058220 to A.B.S, and a UW ADRC Developmental Project (NIH grant P30AG066509) to A.B.S.. M.R.V was supported by an NIH Pathway to Independence Award from NIGMS (1K99GM155552-01), and both M.R.V and S.C.B. were supported by a training grant (T32) from the NIH (2T32GM007454-46). E.S. was supported by a Curci Fellowship, as well as a training grant (T32) from the NIH (T32HG000035). Competing interests: A.B.S., E.G.S., and Y.M. are co-inventors on the U.S. Provisional Patent Application 63/687,924 that includes discoveries described in this manuscript regarding ‘Chromatin Stenciling’™ using DAF-seq.

## Methods

### Bacterial strains and culture conditions

All bacterial strains used in this study were grown in Lysogeny Broth (LB) at 37 °C or on LB medium solidified with agar (RPI, cat# L24030-100.0). Filter sterilized kanamycin (Gold Biotechnology, cat# K-120) (100 mg/L for plasmid propagation, or 30 mg/L for protein expression), and IPTG (ThermoFisher, cat# R0393) (0.5 mM) were added to culture when necessary. *E. coli* strains DH5α (NEB, cat# C2987H), BL21(DE3) (NEB, cat# C2527H) were used for cloning and producing plasmids, and protein expression respectively.

### Cloning and purification of SsDddA

The genes for SsDddA WT, SsDddA5, and corresponding immunity protein (SsDddI), which is required for SsDddA purification, were codon optimized for *E. coli* expression and synthesized as gBlocks with corresponding restriction enzyme recognition sites flanking each end by IDT. The SsDddI was inserted between NdeI and XhoI, and WT SsDddA or SsDddA5 was inserted between NcoI and NotI with a N-terminal 6xHis tag of the vector pColADuet-1 (LifescienceMarket, #PVT0105). The deaminases were cloned into the vector after the immunity protein was successfully cloned into the vector. The whole plasmid sequence was confirmed by PlasmidSaurus.

The purification of deaminases was performed as previously described^6^ with the following modifications: protein expression was induced at 16 °C overnight, cells were resuspended in Ni-NTA Buffer A (50 mM Tris-HCl, pH 7.8, 600 mL NaCl, 10% Glycerol, 10 mM 2-Mercaptoethanol, 0.1% Triton-100) with protease inhibitor cocktail (Thermo Scientific, cat#PIA32955). The cell lysate was loaded on a HisTrap HP His tag protein purification column (5 mL, Cytiva # 17524801) and the deaminase-immunity protein complex was eluted during a gradient from 100% Ni-NTA Buffer A to 100% Ni-NTA Buffer B (50 mM Tris-HCl, pH 8, 600 mL NaCl, 500 mM imidazole, 10% Glycerol) using the NGC Quest 10 Plus Chromatography System (Bio-Rad, #7880003) and the fractions of corresponding A280 peaks were pooled. The major peak eluted during the gradient was pooled, verified using SDS-PAGE, and collected for the denaturing and renaturing steps to separate the immunity protein and the deaminase. The pooled protein complex samples were added to denaturing buffer (50 mM Tris-HCl pH 7.8, 20 mM imidazole, 500 mM NaCl, 6 M guanidine HCl, prepared from Guanidine-HCl (ThermoFisher, cat# 24110), and 5 mM 2-Mercaptoethanol) at 1:10 (v:v) ratio and incubated overnight. The mixture was then loaded back to the 5 mL HisTrap column, with 50 mL of denaturing buffer. The deaminase was renatured during a gradient from 100% denaturing buffer to 100% renaturing buffer (50 mM Tris-HCl pH 7.8, 500 mM NaCl, 10 µM ZnCl_2_, and 10 mM 2-Mercaptoethanol) at 1 ml/min and washed with additional 50 mL renaturing buffer. The renatured deaminase was further purified using HiLoad Superdex 200 pg preparative SEC column (120 mL, Cytiva # 28989335) with storage buffer (50 mM Tris-HCl pH 7.8, 500 mM NaCl, 10 µM ZnCl_2_, 1 mM DTT, and 10% Glycerol). The fractions were evaluated by SDS-PAGE, and the purest fractions were pooled, aliquot into 20 µL stocks, flash frozen with liquid nitrogen, and stored at -80 °C. All Tris-based buffers were prepared from 1 M Tris-HCl (pH8, molecular biology grade ultrapure, ThermoScientific, cat# J22638-K2).

### Mass Spectrometry validation of SsDddA activity (and inactivation)

SsDddA activity was validated via quantification of deoxycytidine deamination by UHPLC-MS/MS. Samples for quantification were treated as previously described with modifications (Kong et al. 2022). In brief, 2 ng/mL of stable isotope-labeled 2-deoxycytidine triphosphate (MilliporeSigma, cat# 646229) and 2-deoxyadenosine triphosphate (MilliporeSigma, cat# 646237) were used as references and added to 50 ng of DNA from each sample before any mass spectrometry sample preparation. The DNA from each sample was mixed with 0.02 U phosphodiesterase I (Worthington, cat# LS003926), 1 U Benzonase (MilliporeSigma, cat# E1014), and 2 U Quick CIP (NEB, cat# M0525S) in digestion buffer (10 mM Tris, 1 mM MgCl, pH 8 at RT) for 3 hours at 37°C, with a total reaction volume of 50 µL. Single nucleotides were separated from the enzymes by collecting the flow-through of a Nanosep centrifugal filter (MWCO 3 kDa, Pall, cat# OD003C33). The UHPLC-MS/MS analysis of cytosine and adenosine was performed on an ACQUITY Premier UPLC System coupled with a XEVO-TQ-XS triple quadrupole mass spectrometer. UPLC was performed on a ZORBAX Eclipse Plus C18 column (2.1 × 50 mm I.D., 1.8 μm particle size) (Agilent, cat# 959757-902) using solvent A consisting of 0.1% ammonium hydroxide in 100% acetonitrile (v/v) and solvent B consisting of 0.1 M ammonium acetate in water, with the following gradient at 0.3 mL/min: 0-1 min 100% A, 1-6 min 100-30% A and 0-70% B, 6-7 min 30-5% A and 70-95% B, 7-8 min 5-100% A and 95-0% B, 8-10 min 100% A. MS/MS analysis was operated in positive ionization mode with 3000 V capillary voltage as well as 350°C and 1000 L/hour nitrogen drying gas. A multiple reaction monitoring (MRM) mode was adopted with the following m/z transition: 227.9 -> 94.82, 227.9 -> 98.98, 227.9 -> 111.98, 227.9 -> 116.99 for dC (collision energy, 32, 18, 6, 12 eV respectively); 238.9 -> 100.8, and 238.9 -> 118.9 for isotope-labeled deoxycytidine (collision energy 32 and 6 eV respectively); 252.10 -> 136.09 for dA (collision energy, 14 eV), 267.1 -> 146.1 for isotope-labeled dA (collision energy, 14 eV) was monitored as well as control. MassLynX was used to quantify the data. All reagents used for mass spectrometer analysis are molecular grade level or above

### Bulk Whole-Genome Amplification (WGA) DAF-seq

2 million K562, HG002, or GM12878 cells were permeabilized as previously described ^2^ with the difference of permeabilized cells being resuspended in freshly prepared Buffer C (15 mM Tris, pH 8.0; 15 mM NaCl; 60 mM KCl; 1mM EDTA, pH 8.0; 0.5 mM EGTA, pH 8.0; 0.5 mM Spermidine, 10 nM ZnCl_2_). The permeabilized cells were treated with 0.25 µM WT SsDddA or SsDddA5 at 25 °C for 10 min. The reaction was quenched with 5% SDS (ThermoFisher, cat# AM9820) before gDNA extraction using HMW DNA Extraction kit (Promega, cat# A2920). The gDNA was then subjected to whole genome amplification with REPLI-G Mini kit (Qiagen, cat#150023) according to the manufacturer’s protocol before being prepared for sequencing.

### Nuclei isolation

GM12878, COLO829BL, and COLO829T cell lines were permeabilized using a digitonin containing isotonic buffer. Briefly, we added 800,000-1,000,000 cells per sample to a 1.5 mL tube (Eppendorf, 022363204) and centrifuged at 400g for 5 min at 4°C. The supernatant was removed, and the cell pellet was resuspended in 100 µL of chilled isotonic Perm Buffer (20 mM Tris-HCl pH 7.4, 150 mM NaCl, 3 mM MgCl2, 0.05% digitonin) by pipette-mixing 10 times. Cells were incubated on ice for 5 min, after which they were diluted with 1 mL of isotonic Wash Buffer (20 mM Tris-HCl pH 7.4, 150 mM NaCl, 3 mM MgCl2, 10 nM ZnCl2) by pipette-mixing five times. Cells were centrifuged at 400g for 5 min at 4°C and the supernatant was removed. The cell pellet was resuspended in chilled isotonic Wash Buffer. Cells were counted using a Cellometer Spectrum Cell Counter (Nexcelom) using ViaStain acridine orange/propidium iodide solution (Nexcelom, C52-0106-5).

### Tissue Preparation

Liver, heart and colon tissue were separately homogenized using a Dounce homogenizer with ten strokes using pestle A, followed by ten strokes using pestle B. 1.5 mL of cold homogenization buffer was added to the sample, which was then filtered through a 70-micron filter. Cells were counted using a Cellometer Spectrum Cell Counter (Nexcelom) using ViaStain acridine orange/propidium iodide solution (Nexcelom, C52-0106-5). 200,000 cells were added to a 1.5 mL tube (Eppendorf, 022363204) and centrifuged at 500g for 5 min at 4°C. The supernatant was removed and the cell pellet was resuspended in 60 µL of chilled Buffer A w/ ZnCl_2_ (15 mM Tris, pH 8.0; 15 mM NaCl; 60 mM KCl; 1mM EDTA, pH 8.0; 0.5 mM EGTA, pH 8.0; 0.5 mM Spermidine, 10 nM ZnCl_2_) by pipette-mixing, followed by addition of 60 ul of 2X lysis buffer (0.025% IGEPAL, 15 mM Tris, pH 8.0; 15 mM NaCl; 60 mM KCl; 1mM EDTA, pH 8.0; 0.5 mM EGTA, pH 8.0; 0.5 mM Spermidine, 10 nM ZnCl_2_). Cells were incubated on ice for 10 minutes and nuclei were counted using a Cellometer Spectrum Cell Counter (Nexcelom) using ViaStain acridine orange/propidium iodide solution (Nexcelom, C52-0106-5). ∼100,000 nuclei were used as input to the SsDddA reaction.

### DddA Reactions

All targeted DAF-seq reactions utilized SsDddA WT. Enzyme optimization experiments targeted *NAPA* and *WASF1 and* treated 100,000 GM12878 permeabilized cells at 25°C for 10 minutes and 20 minutes, at enzyme concentrations of 0.25 μM, 1 μM, and 4 μM. All subsequent SsDddA reactions were performed at 25°C for 10 minutes with a 4 μM enzyme concentration. Reactions were neutralized by the addition of 20% sodium dodecyl sulfate (SDS) to a final concentration of 5%. Genomic DNA was extracted using the Monarch Genomic DNA Purification Kit (New England Biolabs, T3010S) and quantified using a Qubit 1X dsDNA High-Sensitivity kit (Invitrogen, Q33231).

### PCR Amplification of SsDddA-treated genomic DNA

Regions-of-interest were amplified from SsDddA-treated genomic DNA in 50 ul reactions consisting of 25 μL LongAmp Hot Start Taq 2X Master Mix (New England Biolabs, M0533L), 2 μL each of 10 μM forward and reverse primers (**Extended Data Table 1**), and 30-100 ng of gDNA. PCR conditions: 94°C for 60s, 35 cycles of 94°C for 30s, 30s annealing, and 65°C extension, followed by a final extension of 65°C for 10 min. Annealing temperatures and extension times varied by target (**Extended Data Table 1)**. Amplicons were purified using the Monarch PCR & DNA Cleanup Kit (New England Biolabs, T1030L).

### Library preparation and sequencing

Libraries were prepared as previously described^26^. Multiplexed library preparation was performed using the SMRTbell prep kit 3.0 (PacBio, cat#102-141-700) and SMRTbell barcoded adapter plate 3.0 (PacBio, cat#102-009-200). Final sequencing libraries were sequenced on the PacBio Revio platform using v3.2 chemistry.

### DAF-seq alignment and preprocessing

PacBio HiFi reads were converted to FastQ format using Samtools fastq^33^ (v1.17, parameters: -T) and aligned to hg38 using minimap2 (v2.22-r1101, parameters: --MD -Y -y -a -x map-pb). C-to-T and G-to-A changes were identified by comparing the sequencing read and the hg38 reference using pysam (v0.21.0, https://github.com/pysam-developers/pysam). Secondary and supplementary alignments were filtered out. The designation of the original SsDddA modified DNA strand as “top” or “bottom” was identified by quantifying the proportion of C-to-T and G-to-A changes. Reads with at least 90% of these changes were classified as “CT” or “GA” and retained for subsequent analyses. C-to-T and G-to-A changes were converted to the IUPAC DNA ambiguity codes Y and R in CT and GA reads, respectively. Modified reads were realigned to hg38 using the same parameters. Identification of modification sensitive patches (MSPs) and the generation of nucleosome and accessibility pileups was done using Fibertools^26^ add-nucleosomes (v0.5.4, parameters: -n 60, -c 70, --min-distance-added 15, -d 10). MSPs > 150bp in length were identified as actuated elements.

### DAF-seq Target Enrichment

The proportion of total primary read alignments that mapped within the target region was calculated for both targeted DAF-seq and Fiber-seq data from the same cell line (GM12878). DAF-seq target enrichment was calculated as the proportion of on-target reads in DAF-seq over Fiber-seq.

### PCR Duplicate Identification

DAF-seq PCR duplicate reads were identified by comparing the deamination status of every position susceptible to deamination (each C on top strand reads, each G on bottom strand reads). This combination of deamination statuses was treated as a unique identifier and all reads sharing this identifier were grouped. One read from each group was randomly selected to be marked as unique while the remaining reads in each group were marked as duplicates.

### CpG Methylation Analysis

NAPA and UBA1 regions were amplified for 30 PCR cycles as described above (**Extended Data Table 1**) using 70 ng of HEK293 genomic DNA as template. 1 μg of amplicons from each target was treated with M.SssI CpG methyltransferase (New England Biolabs, M0226S) per the vendors recommendations. 600 ng of M.SssI treated and untreated amplicons were treated with SsDddA for 10 minutes at 25°C. DNA treated with SsDddA and M.SssI, and M.SssI only (5mC negative control) were re-amplified for 20 PCR cycles. These two samples along with M.SssI treated DNA (5mC positive control) were sequenced using PacBio HiFi as described above. Amplicons were purified after each PCR using the Monarch PCR & DNA Cleanup Kit (New England Biolabs, T1030L).

Sequencing reads from each target were aligned to reference fasta files containing only the target region. Primary alignments from the M.SssI + SsDddA treatment beginning and ending within 100bp of the region boundaries were used in analysis. Genomic positions were grouped into CpG and non-CpG cytosine (or guanine for bottom strand) categories, and the deamination occurrences at each position within each read were aggregated. Positions within 28bp of the target region boundaries overlapped priming sites and were omitted from analysis. Positions CpG methylation of the 5mC positive control was quantified using pb-CpG-tools (aligned_bam_to_cpg_scores, v2.3.1, https://github.com/PacificBiosciences/pb-CpG-tools).

### Deamination Motif Analysis

Sequence bias in SsDddA activity was evaluated for GM12878 WGA realigned data using pysam (v0.21.0, https://github.com/pysam-developers/pysam), considering only primary alignments. For each deaminated base, denoted in the read sequence by the ambiguity codes Y and R, the seven base (7mer) reference sequence centered on the deaminated base was tracked. Sequences originating from GA reads were reverse-complemented to orient the 7mer to the cytosine context. All 7mers were combined to generate a position weight matrix (PWM) which was used to generate a sequence logo (Logomaker v0.8)^34^.

### Transcription Factor Footprinting and Codependency

We used FIMO^35^ to scan the hg38 reference sequence of an applicable region for transcription factor motifs contained within the JASPAR database ^36^ and filtered the results for motifs with a q-value <= 0.05. We then used Fibertools^26^ footprint to identify single-fiber TF footprints for each motif. We filtered motifs to those footprinted on >= 5% of fibers on both top (CT) and bottom (GA) strands. Footprinted motifs that overlapped either motif by 80% were merged using Bedops^37^ (v2.4.41), and the resulting elements that overlapped by 90% were merged to combine elements encompassed entirely within a larger element. We quantified occupancy within each merged element as the proportion of fibers footprinted at a motif contained within the merged element. TF co-occupancy and codependency were calculated as described previously^12^. Briefly, for each pair of footprinted elements, we calculated the expected co-occupancy as the product of their proportion of fibers bound, while the observed co-occupancy was calculated as the proportion of fibers with an accessible element (MSP > 150 bp) spanning both elements and bound at each element. We quantified the essentiality of each element by constructing codependency graphs, with nodes representing TF elements and edge weights representing codependency scores. We constructed graphs that omitted individual elements and limited each analysis to fibers accessible but unbound at that respective element. We also constructed a baseline codependency graph containing all elements. We quantified the total codependency of each graph by summing all edge weights and normalizing by the number of edges. Finally, we calculated essentiality as the ratio of total codependency of the baseline graph over element-excluded graphs, with a higher ratio indicating a larger reduction in codependency following the loss of TF binding at that element.

### Identification of heterozygous positions

Top and bottom strand base calls were counted at each genomic position within each target region for each primary read alignment. Positions within 25bp of the target region boundaries overlapped priming sites and were omitted from analysis. Top strand and bottom strand base call proportions were plotted in R using the ggplot2 hexbin package (parameters: bins = 30).

### *UBA1* Transcription Start Site Co-actuation and Codependency

*UBA1* bottom strand reads were assigned haplotypes according to base calls at chrX:47,194,331 (rs56269549). The four *UBA1* transcriptional start sites (TSSs) were identified previously using full-length transcript data^16^. MSPs > 150bp long were intersected with each TSS using Bedops^37^ (bedmap, parameters: --ec --fraction-ref 0.8 --echo --echo-map-id, v2.4.41). TSS co-actuation was calculated as the proportion of fibers with actuated elements overlapping both TSSs. Codependency was calculated as above using TSS co-actuation instead of TF co-occupancy.

### DAF-seq Clustering

Liver and GM12878 reads were assigned haplotypes according to base calls at chr8:144,416,180 (rs2280839). Reads from each sample with a hamming distance of 3 or less were identified as duplicates and one read from each group was randomly selected as a unique read. The deamination status at each applicable genomic position was recorded and used as a feature for clustering DAF-seq reads. Clustering analysis was performed using Scanpy (v1.10.3). Briefly, 5000 unique reads from each haplotype of each tissue were randomly selected. A neighborhood graph was computed (pp.neighbors, parameters: n_pcs=0, n_neighbors=200) and visualized with Uniform Manifold Approximation and Projection (UMAP). Clustering was performed using the Leiden algorithm (tl.leiden, parameters: flavor="igraph"). Clusters containing fewer than 1000 fibers were removed. Fibers from cluster 1 were re-clustered using the same parameters. Sub-clusters containing fewer than 200 fibers were removed.

### COLO829 cell mixture

Pure populations of COLO829BL cells (cat# CRL-1980, lot# 70022927) and COLO829 (referred to as COLO829T) cells (cat# CRL-1974, lot# 70024393) were obtained from ATCC and expanded. All cells were grown at 37°C with 5% CO2. COLO829BL lymphoblastic suspension cells were grown in RPMI-1640 (Fisher cat# 11875093) with 10% fetal bovine serum (FBS, Fisher cat# 10082147) shaking at 90 rpm. COLO829T adherent melanoma cells were grown in RPMI-1640 10% FBS. Both cell lines were expanded and cryopreserved in freezing media consisting of RPMI-1640 10% FBS with 10% DMSO (Sigma Aldrich cat# D2650-100ML) at a cell density of 3 million viable cells per mL. Cryopreserved COLO828T and COLO829BL cells were thawed and mixed in a 1:50 ratio using viable cell counts based on Countess cell counter readings (Invitrogen Thermo Fisher) with trypan blue (Fisher cat# 15250061). Cells were aliquoted into cryovials with constant swirling to maintain the homogeneity of the mix. Mixed cells were cryopreserved at a cell density of 2.55 million viable cells per mL.

### Single-cell Fluorescence Activated Cell Sorting (FACS)

DddA-treated cells were sorted on a BD FACSAriaII using sequential gating on forward-scatter (FSC) and side-scatter (FSC) to detect doublets and debris. Individual cells were sorted into wells of a 96-well plate containing 3 ul of Cell Buffer (BioSkryb Genomics, 100183). Cells were immediately flash frozen on dry ice.

### Single-cell PTA Whole Genome Amplification

After dispensation of individual nuclei into independent plate wells, the ResolveServices (SM) team performed the Services Custom PacBio Long Read Amplification (BioSkryb Genomics 101157). Briefly, the ResolveDNA(TM) workflow was customized to increase amplicon size. After 2.5 hours of isothermal amplification, samples were quantified using qubit HS DNA (Thermofisher Q33231) and shipped to the Stergachis laboratory for library preparation and sequencing.

### Single-cell read collapsing

PacBio HiFi reads were converted to FastQ format using Samtools fastq^33^ (v1.17, parameters: -T) and aligned to hg38 using minimap2 (v2.22-r1101, parameters: --MD -Y -y -a -x map-pb). Soft-clipped bases were trimmed from the alignments to remove chimeric portions of each read. The original template strand of clipped reads was identified as “top” or “bottom” strand by quantifying the proportion of C-to-T and G-to-A changes relative to the hg38 reference using pysam (v0.21.0, https://github.com/pysam-developers/pysam). Primary alignments with at least 95% of these changes were classified as “CT” or “GA” and retained for subsequent analyses.

The hg38 reference genome was partitioned into overlapping 150 bp bins with a 25 bp sliding window. For each bin, the sequence hamming distance between reads that completely spanned that bin was computed, ignoring insertion and deletions. Similar reads were identified as those sharing a minimum of 11 bins (400 bp in total) and with >= 99% identical sequence within >= 80% of shared bins. Similar reads were sequentially grouped together such that each read shared similarity with at least one other read within the group.

Consensus sequences were generated for each group as follows: for each reference position represented within the read group, the composition of base calls at that position was quantified. In cases with disagreement between reads, bases comprising >= 50% of all base calls was used, except in cases where the two most common bases are C & T or G & A and comprised >= 50% of all base calls, in which case C or G was used to overcome putative spurious post-amplification deamination. Deletions and insertions were ignored unless they were present in all reads at that position. When insertions were present in every read the shortest insertion was used. Consensus reads were assigned a random identifier and converted to fasta format. Consensus reads were realigned to hg38 as above and C-to-T and G-to-A changes were converted to the IUPAC DNA ambiguity codes Y and R in CT and GA reads, respectively.

### Consensus read phasing and coverage quantification

Consensus reads were assigned haplotypes with Whatshap haplotag^38^ (v2.3, parameters: --ignore-read-groups, --output-haplotag-list) using previously phased parental variants^16^. The mappable hg38 genome was computed as regions not overlapping the ENCODE blacklist (accession ENCFF356LFX)^39^ or hg38 regions with unreliable coverage in HG002 Fiber-seq data^16^. Regions of greater than 400bp contiguous bases that consisted of more than 1 consensus read per haplotype strand in at least one cell were identified using Samtools depth^33^ (v1.17) and excluded from the hg38 mappable genome. Coverage calculations were performed using Samtools depth in combination with Bedtools^40^ (v2.31.0) and Bedops^37^ (v2.4.41). Read length statistics were calculated in python using custom scripts.

### Single-cell chromatin analyses

Fibertools^26^ add-nucleosomes (v0.5.4, parameters: -n 60, -c 70, --min-distance-added 10) was used to calculate deamination autocorrelation and identify modification sensitive patches (MSPs) within consensus reads. HG002 regulatory elements were identified previously from Fiber-seq data using the FIRE pipeline^16^. scDAF-seq regulatory elements were classified as actuated if they were overlapped by an MSP of >150bp on either strand. These overlaps were required to span at least 50% of either the length of the MSP or the length of the FIRE peak. Read lengths, deamination rates, Jaccard distances, and percent actuation were computed in python using custom scripts.

---

**Extended Data Figure 1.**
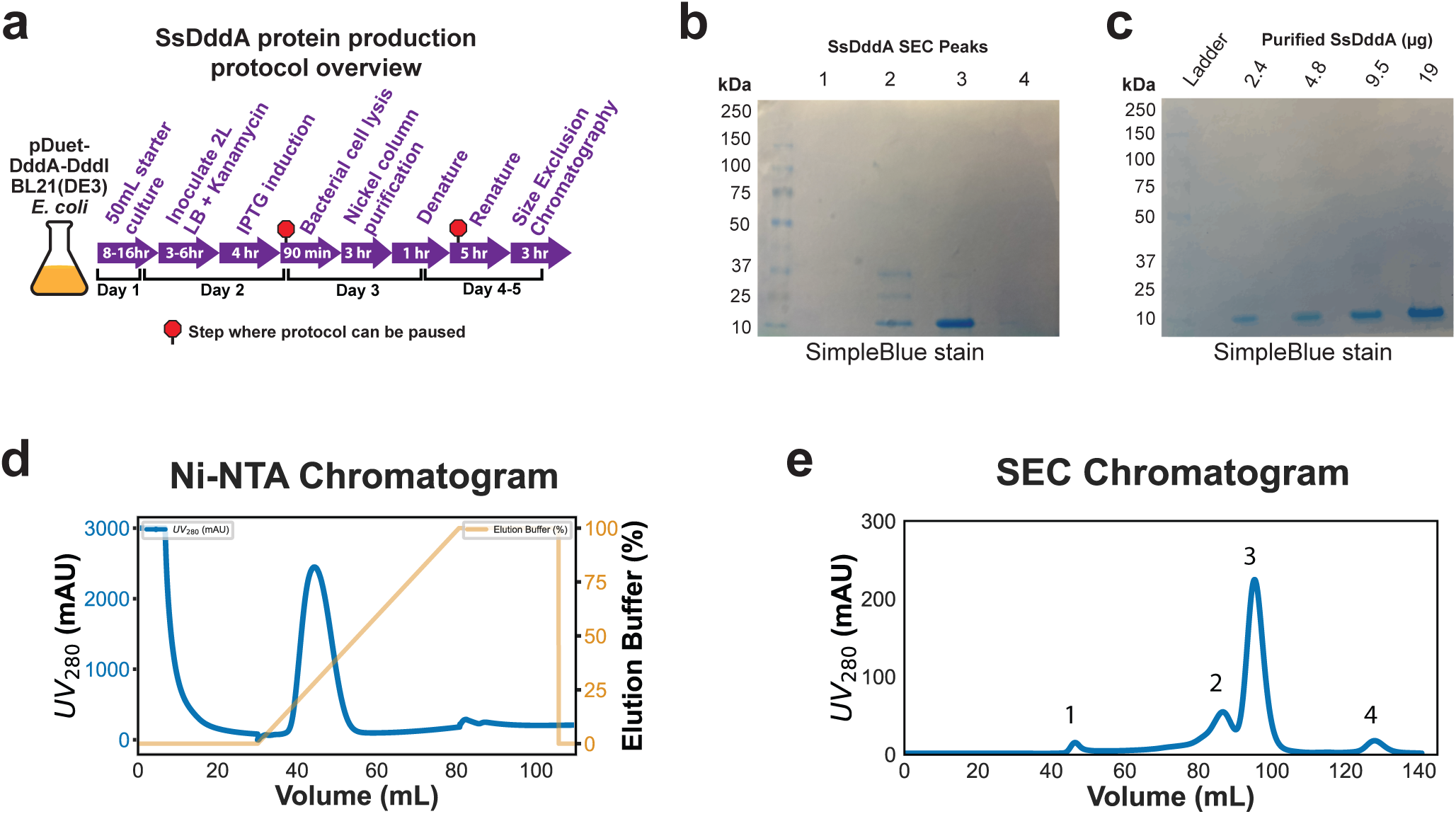
Production of recombinant SsDddA variants. **(a)** Schematic representation of SsDddA production workflow. **(b)** SDS-PAGE analysis of peak fractions collected from Size Exclusion Chromatography (SEC) as indicated in panel (e). **(c)** SDS-PAGE gel depicting the purified SsDddA protein. **(d)** Representative Ni-NTA affinity chromatogram from the Nickel column purification step. **(e)** SEC chromatogram with labeled peaks (1, 2, 3, and 4) corresponding to distinct fractions, pooled and analyzed by SDS-PAGE as shown in panel (c).

**Extended Data Figure 2.**
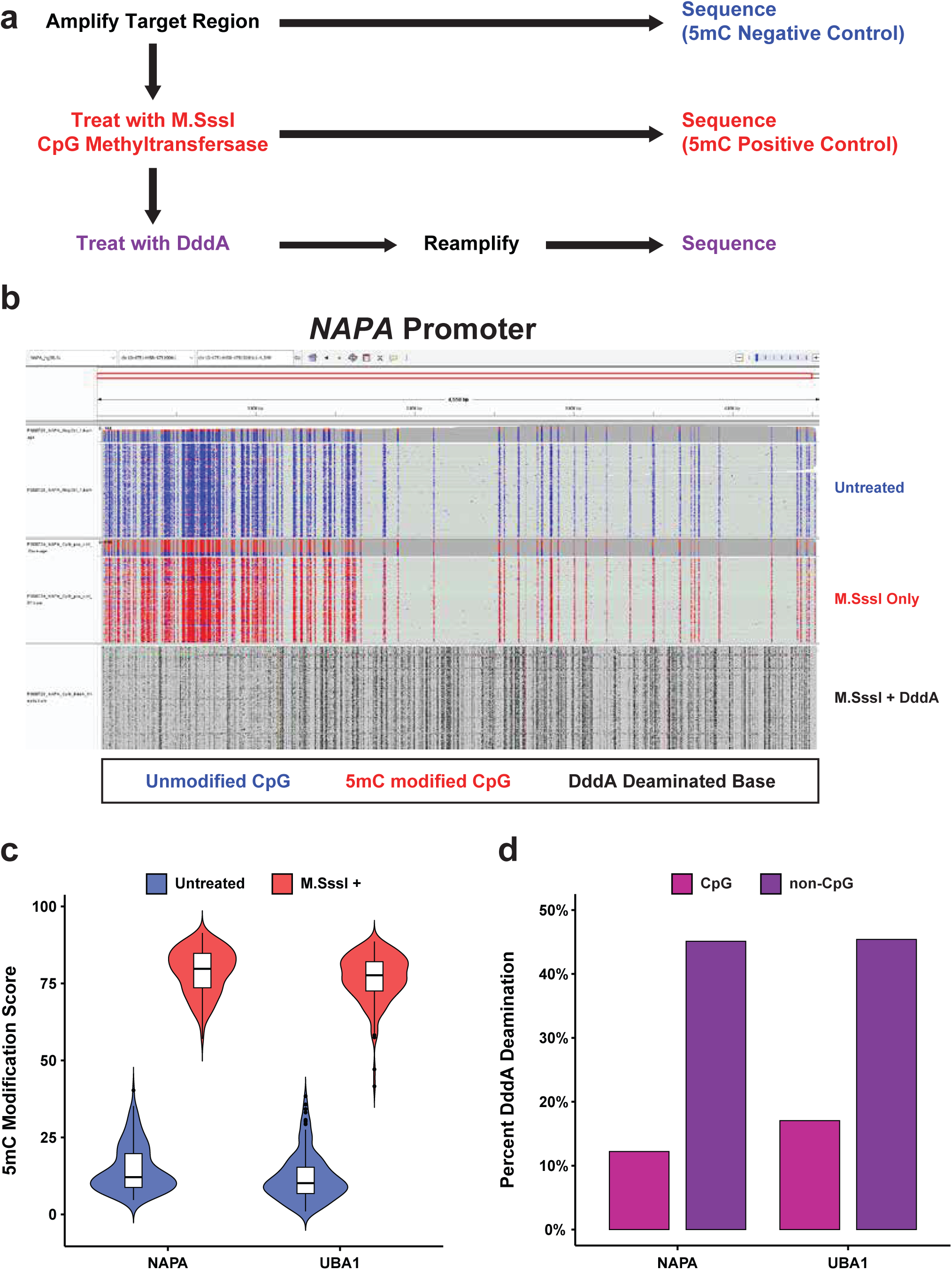
SsDddA activity at 5mCpG dinucleotides. **(a)** Experimental overview of the comparison of SsDddA deamination rate between 5mC methylated and unmethylated cytidines. **(b)** IGV browser displaying CpG methylation for 5mC negative control (top), 5mC positive control (middle), and M.SssI plus SsDddA treated DNA (bottom). **(c)** Violin plots displaying 5mC modification scores at CpG dinucleotides for untreated and M.SssI treated controls. **(d)** Percentage of cytosine deamination by SsDddA grouped by CpG dinucleotides and all other cytosine bases. Data is shown for two targeted regions, the *NAPA* promoter region and the *UBA1* promoter region.

**Extended Data Figure 3.**
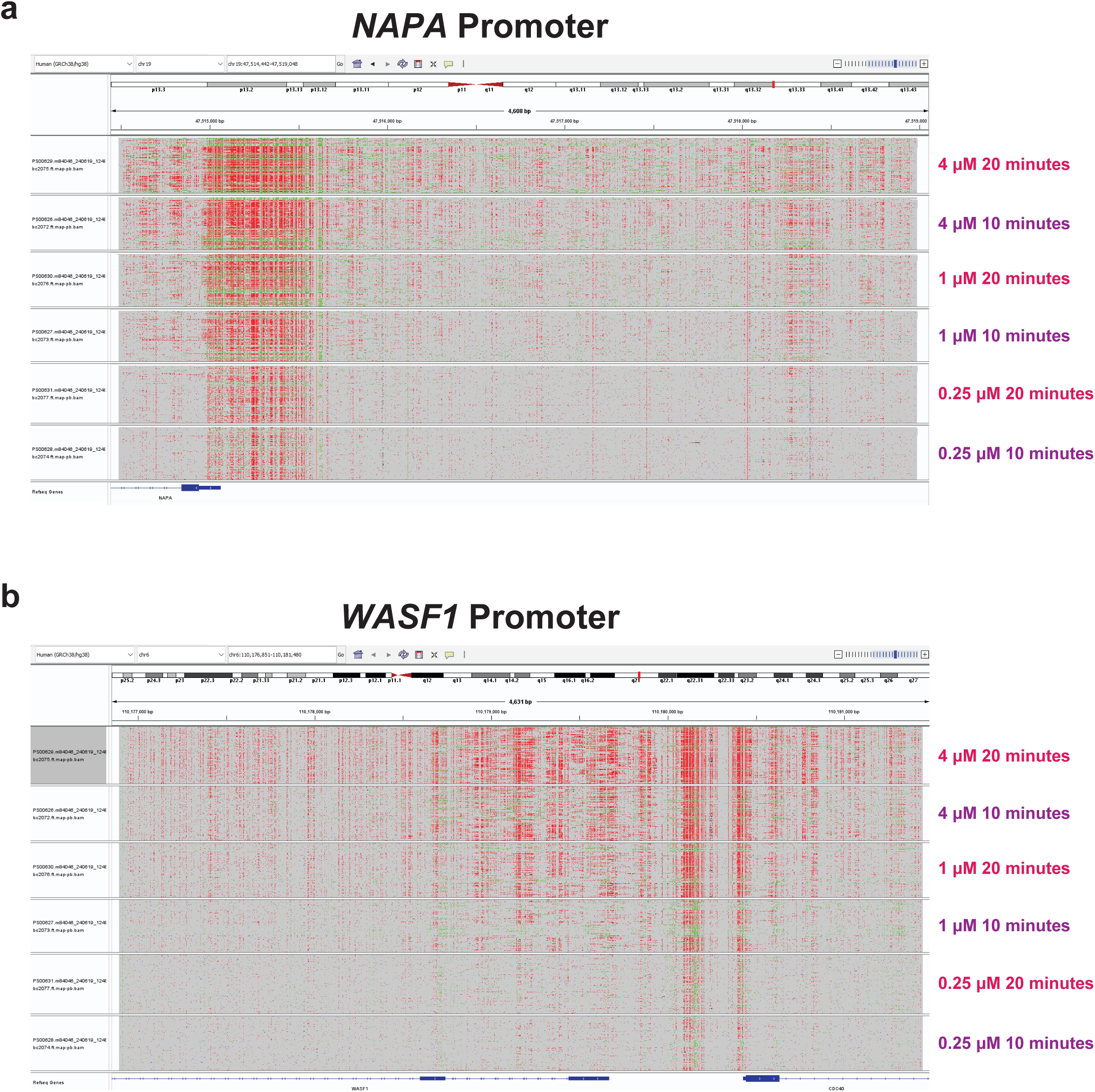
Deamination frequency by SsDddA enzyme concentration. **(a)** IGV browser displaying DAF-seq reads from GM12878 cells treated with a range of SsDddA enzyme concentrations and treatment times for a 4.5 kb region spanning the *NAPA* promoter. **(b)** Same as in a for a 4.5 kb region spanning the *WASF1* promoter.

**Extended Data Figure 4.**
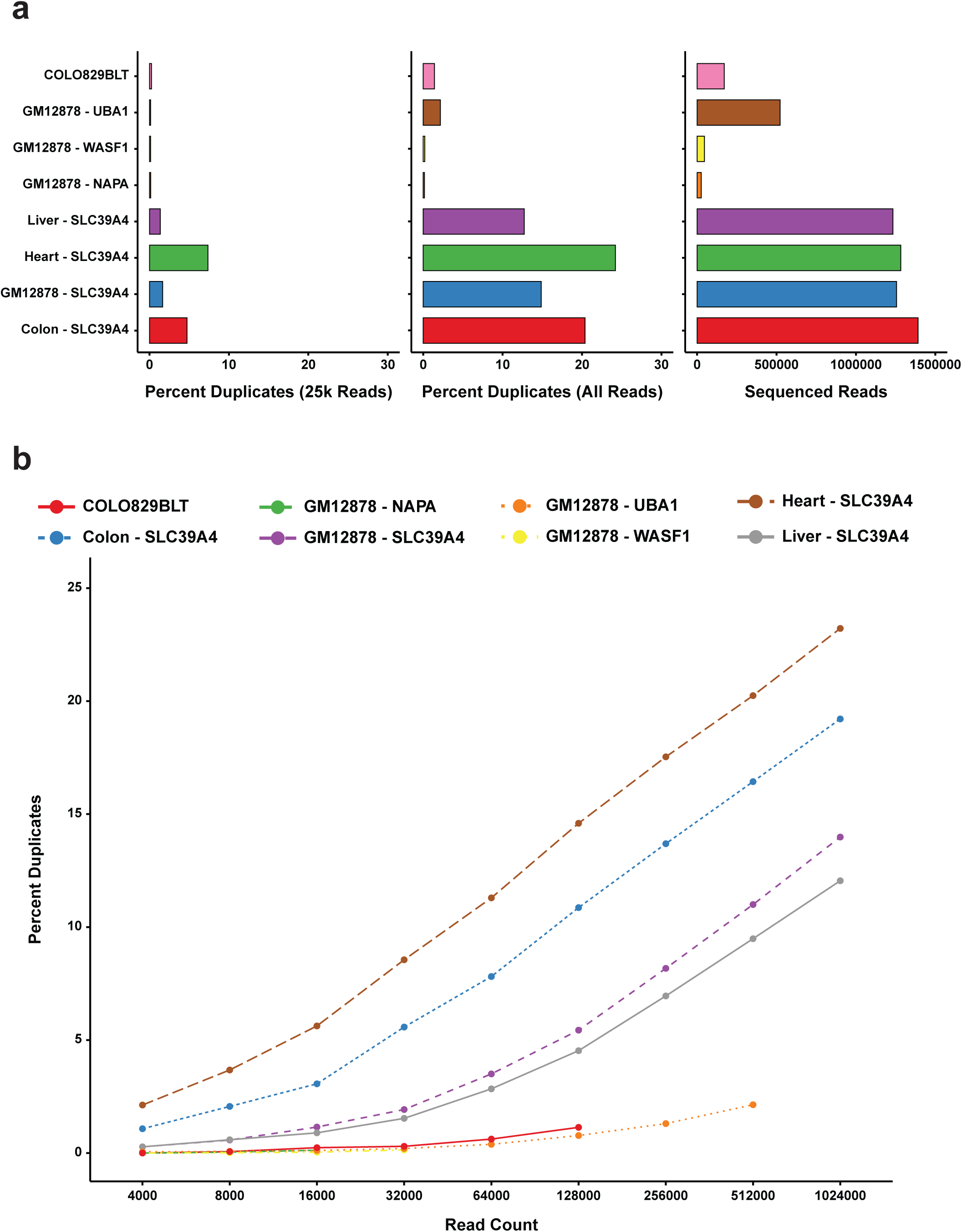
Identification of PCR duplicates via chromatin architectures. **(a)** Barplots for targeted DAF-seq libraries displaying the percentage of reads with identical deamination profiles (duplicate reads) at a downsampled depth of 25,000 reads (left) and all sequenced reads (middle), and the total number of sequenced reads (right). **(b)** Duplication rates for each of the libraries in **a** at different downsampled read counts.

**Extended Data Figure 5.**
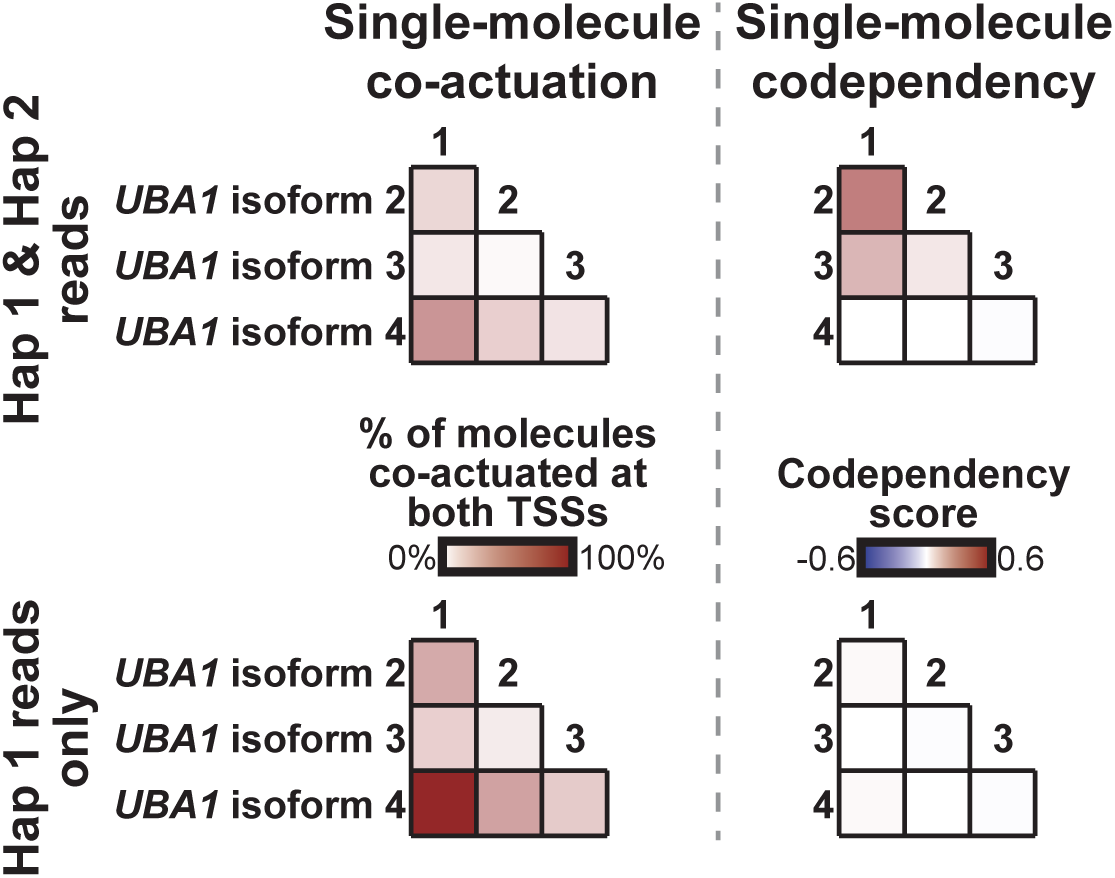
*UBA1* transcriptional start site codependency. Co-actuation (left) at *UBA1* transcriptional start sites (TSSs) for all reads regardless of haplotype (top) and haplotype 1 reads only (bottom). *UBA1* TSS codependency (right) for all reads regardless of haplotype (top) and haplotype 1 reads only (bottom).

**Extended Data Figure 6.**
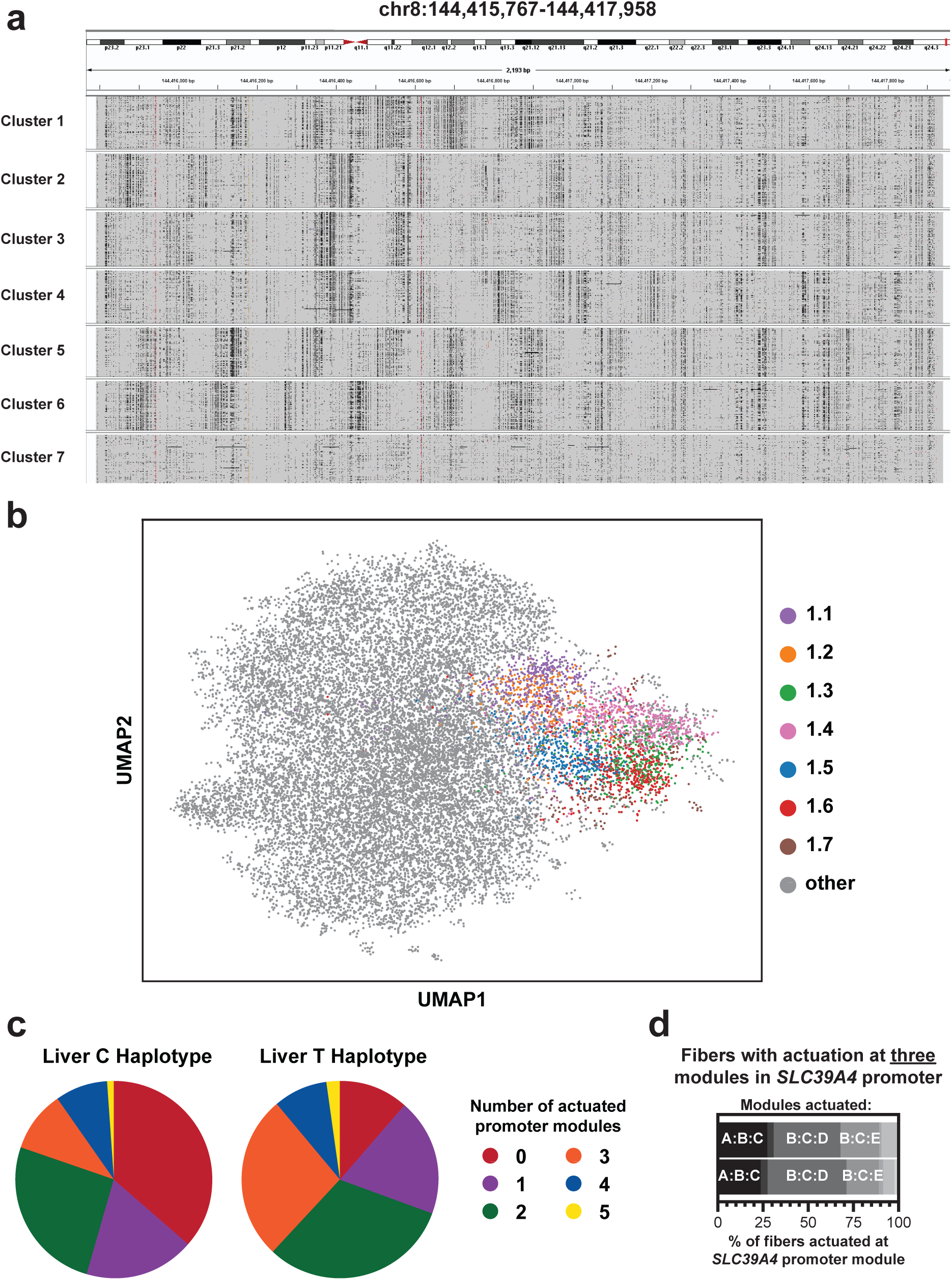
Clustering of *SLC39A4* chromatin architectures. **(a)** IGV browser displaying representative DAF-seq reads from each *SLC39A4* cluster. Black tick marks represent SsDddA deamination positions. **(b)** UMAP from Fig. 4b highlighting fibers from each sub-cluster in Fig. 4d. **(c)** Pie charts displaying the number of liver fibers from cluster 1 that are actuated at each count of actuated *SLC39A4* promoter modules described in Fig. 4e. **(d)** Stacked barplots quantifying combinatorial module usage within liver fibers actuated at three modules.

**Extended Data Figure 7.**
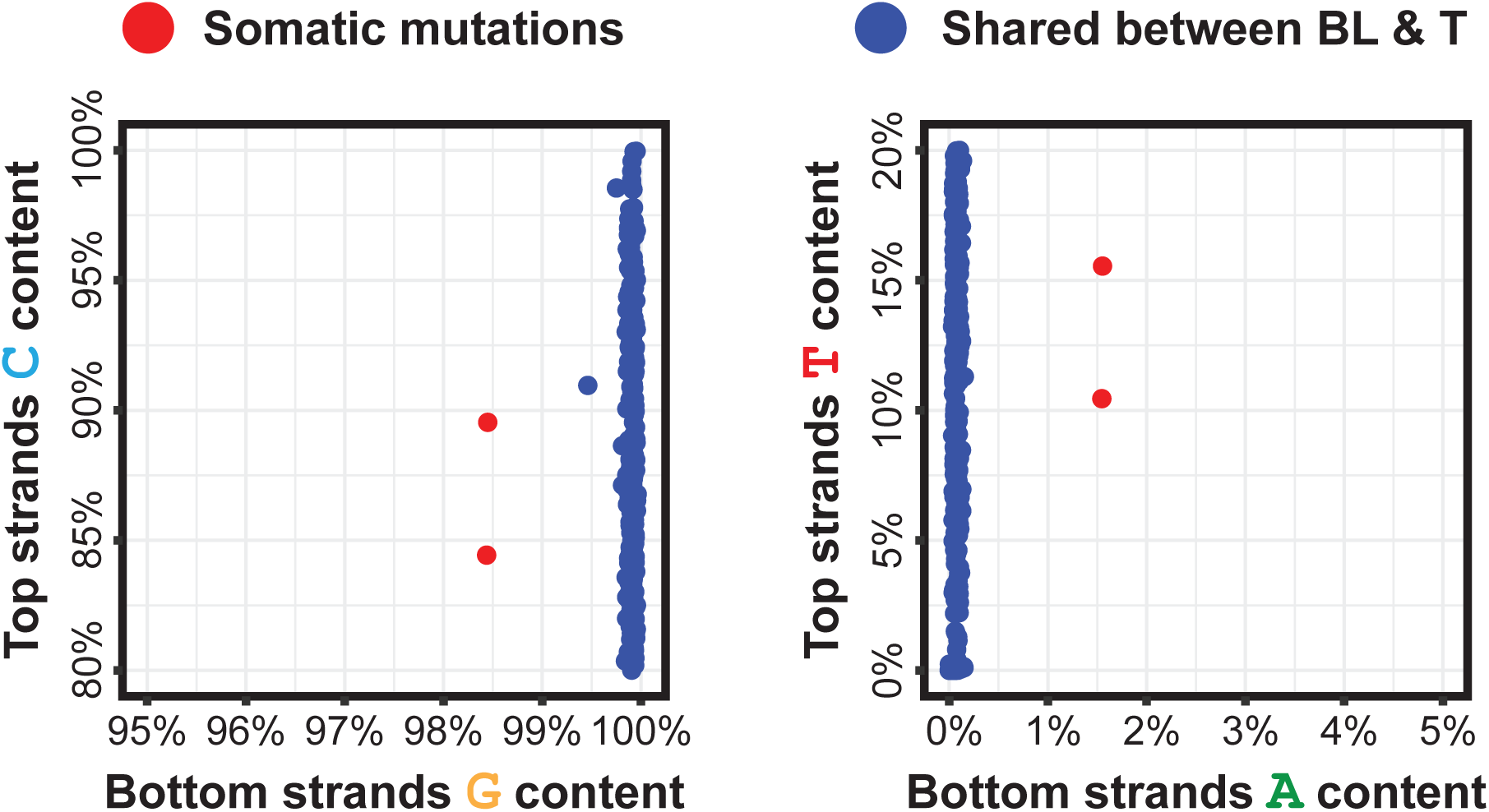
Identification of low-frequency genetic variants. Hexbin plots showing the targeted DAF-seq base content at each genomic position in a 3.8 kb region described in Fig. 5. The two positions comprising the somatic dinucleotide mutation CC>TT on one haplotype of COLO829T (chr17:19,447,245-19,447,246) are colored red. All other positions are colored blue.

**Extended Data Figure 8.**
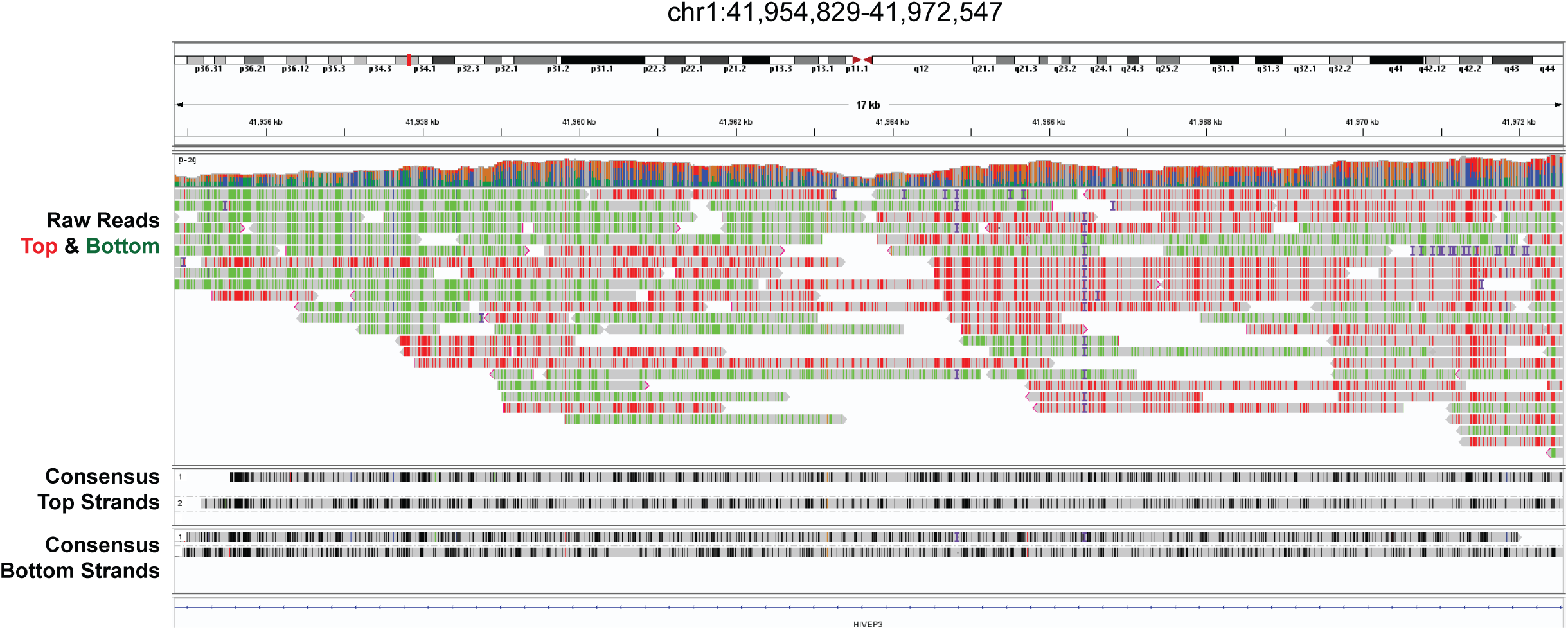
Generation of single-cell consensus reads. IGV browser displaying single-cell DAF-seq reads from one cell. Deamination events for raw reads (top) are colored red for reads originating from the top strand and green for reads originating from the bottom strand. Consensus reads (bottom) are grouped by strand (top or bottom) and ordered by haplotype.

**Extended Data Figure 9.**
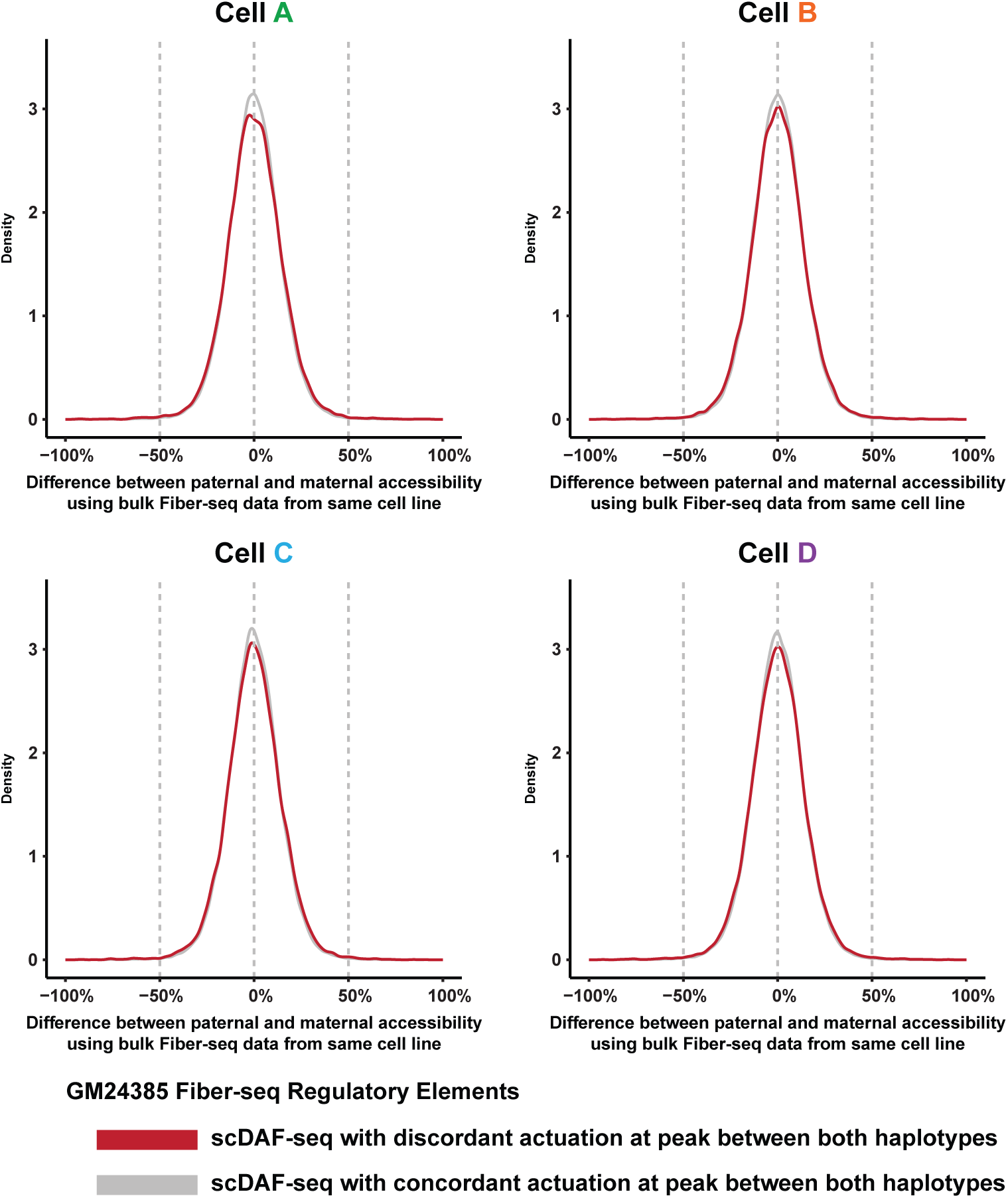
Haplotype selectivity of discordant regulatory elements. Density plots for each cell displaying Fiber-seq chromatin actuation of GM24385 regulatory elements by haplotype. Regulatory elements that are discordant by haplotype within that cell are shown in red. All other regulatory elements (not discordant by haplotype within that cell) are shown in gray.

